# De novo assembly of the *Trypanosoma congolense* genome reveals an organisation influenced by antigenic variation but distinct from *Trypanosoma brucei*

**DOI:** 10.64898/2026.02.19.706783

**Authors:** Marija Krasiļņikova, Jane C Munday, Dario Beraldi, Stephen Larcombe, Guy Oldrieve, Craig Lapsley, Liam J Morrison, Keith Matthews, Richard McCulloch

## Abstract

Antigenic variation is a widespread process for pathogen evasion of mammalian adaptive immunity, involving the continuous change in exposed antigens. In African trypanosomes, antigenic variation involves changes in the expression of Variant Surface Glycoprotein (VSG). Understanding of VSG expression control and change amongst African trypanosomes is most advanced in *Trypanosoma brucei*. In the important animal trypanosome, *Trypanosoma congolense*, incomplete assembly of the genome has held back understanding of the mechanics of antigenic variation. Here, we have used long-read DNA sequencing and Hi-C DNA interaction analysis to provide a telomere-telomere assembly of the *T. congolense* genome. This assembly reveals a genome comprising 12 diploid chromosomes, one tetraploid chromosome, and more than 100 small chromosomes. Within this new assembly, we reveal several features of *VSG* organisation and expression that differ from *T. brucei*. The majority of the *T. congolense VSG* archive, estimated at ∼1500 genes, localises to subtelomeres in 12 of the 13 large chromosomes, but these loci are notably smaller than are found in *T. brucei*. Furthermore, transcriptome analysis reveals expression of *VSG*s across the *T. congolense* subtelomeres, which are not separated within the nucleus from non-*VSG* chromosome regions, indicating that there is no dedicated VSG expression site. Strikingly, one chromosome contains approximately 40% of the *VSG* archive and is largely transcriptionally silent, potentially acting as the major reservoir of new VSG variants. Finally, we show that *VSG* expression can be detected from multiple small chromosomes. In summary, the new genome assembly provides a platform for understanding a potentially unusual operation of VSG expression and switching in *T. congolense*.

## Introduction

African trypanosomes, including *Trypanosoma brucei*, *Trypanosoma congolense* and *Trypanosoma vivax*, rely on antigenic variation to evade elimination by the adaptive immune response of mammals^1^. In each case, antigenic variation is thought to involve periodic changes in expression of Variant Surface Glycoprotein (VSG)^2,3^. Our understanding of this critical survival mechanisms is most advanced in *T. brucei*^4,5^, whose genome has been shaped by the need to express one VSG in a cell at a time^6–8^ and to house a store of silent *VSG* genes that can be expressed in the course of an infection^9,10^. Expression of one VSG/cell in the mammal is dictated by selective transcription of a single *VSG* from one of ∼15 bloodstream *VSG* expression sites (BESs), all of which are found abutting the telomeric ends of many *T. brucei* chromosomes^11^. Most of the *T. brucei* genome is found on 11 megabase chromosomes (∼1-5 Mbp size range), each of which is separated into two compartments: a stable, diploid core that is unusually heavily transcribed due to virtually all genes being expressed from multigenic transcription units^12^; and less stable, transcriptionally silent subtelomeres that separate the chromosome cores from telomeric *VSG* BESs^9^. The subtelomere compartment contains arrays of ∼2000 silent *VSG*s and *VSG* pseudogenes^13,14^ and its total size approaches that of the core^15,16^. Moreover, subtelomeres are notably variable between *T. brucei* strains^17^, between chromosome homologues^18^, and over time in culture, in particular after mutation of the homologous recombination (HR) factors RAD51 or BRCA2^16,19^. Such variability appears consistent with Hi-C evidence for greater interactivity of chromosome subtelomeres relative to the cores^15^. At least part of this subtelomere variability likely stems from recombination of silent *VSG*s into BES to change the expressed VSG during antigenic variation. Such recombination is not limited to the *VSG* archive in the megabase chromosomes, since *T. brucei* also possesses many sub-megabase chromosomes, often named mini- (∼50-150 kbp in size) or intermediate-chromosomes (∼150-700 kbp). Telomere-proximal *VSG* BES and silent *VSG*s are found on many of these chromosomes, thus expanding the *VSG* archive^20^, though some also contain other genes, which are expressed^16^. A shared feature of sub-megabase chromosomes is 177 bp repeats^20^, which appear to act as origins of DNA replication^16^ and recruit many kinetochore components (Carloni et al Biorxiv 2025.04.22.649942v1).

Our understanding of the organisation of the *T. congolense* genome and how it is influenced by antigenic variation is substantially less advanced than that of *T. brucei*. Fewer pulsed field gel analyses of *T. congolense* have been conducted than for *T. brucei*, meaning the total number of *T. congolense* chromosomes is not known, though there is evidence for numerous submegabase chromosomes^21–24^. Though recombination of *T. congolense VSG*s has been linked to expression^25^, it is unclear if such recombination is the driver of immune evasion. Indeed, expression of a *VSG* from a minichromosome has been documented without recombination^26^. Such a finding is consistent with long-read PacBio sequencing that revealed potential BES facsimiles on several *T. congolense* minichromosomes (which lack 177 bp repeats)^23^. However, though such putative minichromosomal BESs comprise telomeric *VSG*s and some upstream conserved sequences^23^, they are unlike *T. brucei* BESs in that they lack detectable expression site associated genes (ESAGs), RNA Pol I promoters or 70 bp repeats^11^, and no work has documented *VSG* transcription from them. More widely, the organisation of the *VSG* archive^27^ in the *T. congolense* genome is unknown, adding to uncertainty about the operation of antigenic variation. It is clear that *T. congolense* has the capacity to encode many VSGs but in contrast to *T. brucei*, where VSGs are separated into a- and b-types (based on cysteine residue distribution^28^) that undergo seemingly unconstrained recombination, *T. congolense* contains only b-type VSGs^2^ that fall into multiple defined phylotypes, with recombination considered to be largely limited to within a phylotype^27,29^. Whether such a constraint on *VSG* recombination is based on *VSG* gene sequence or aspects of genome location or transcription is unknown, since the locations of *VSG*s across the full *T. congolense* genome has not been documented. In this regard, no work has documented sequences around the telomeres of all chromosomes, and so it remains unclear if canonical *VSG* BESs are present in the *T. congolense* genome but have escaped detection.

Here, we have addressed many of these questions around how antigenic variation has shaped the *T. congolense* genome by using long-read Oxford Nanopore Technology (ONT) sequencing allied to Hi-C to generate a *de novo* assembly of the *T. congolense* IL3000 genome. Using this approach, we show that the *T. congolense* genome comprises 13 ‘megabase’ chromosomes, all of which were resolved as haplotypes and assembled telomere-telomere. Many more submegabase chromosomes were similarly fully assembled. 12 of the 13 megabase chromosomes are diploid, while one is tetraploid in wild type cells but reduces in ploidy in the absence of the HR factor RAD51. We show that the organisation of *VSG*s in the genome is quite different from that of *T. brucei*: VSG-rich subtelomeres in *T. congolense* megabase chromosomes are notably smaller than in *T. brucei*, are not transcriptionally silent, and do not show Hi-C evidence for being a distinct compartment to the core. Furthermore, though a silent *VSG* archive is present, it apperas limited to a single chromosome pair that is largely untranscribed and whose ploidy is reduced after loss of BRCA2 or RAD51.

## Results and discussion

### Assembly of the *T. congolense* IL3000 genome

To attempt to assemble the *T. congolense* IL3000 genome, we prepared DNA from wild type, RAD51 null mutant and BRCA2 null mutant bloodstream cells (Fig.S1) growing in culture and compared several sequencing and assembly approaches (Fig.S2). Based on a combination of genome completeness scores determined by QV^30^ and BUSCO^31^, as well as contiguity of the haplotype-resolved sequences, the most complete assembly of the largest chromosomes was generated by verkko^32^, combining ONT long read data and Hi-C reads. In contrast, the more numerous smaller chromosomes were best assembled using hifiasm^33^. Thus, a combination of the two assemblies provided a description of the whole nuclear genome. The resulting combined assembly, following gap closure, comprised 28 contigs >850 kbp, all of which displayed telomeric repeats at each contig end (TTAGGG/CCCTAA) (Fig.1A). In addition, two intermediate-sized telomere-telomere contigs (236,893 bp and 415,598 bp) and 145 telomere-telomere <100 kbp contigs were also assembled, with both of these sets of sequences containing 369 bp repeats previously shown to be characteristic of the small chromosomes of this *Trypanosoma* species^23^. Whether or not the 369 bp repeats provide the same DNA replication origin^16^ and putative centromere functions (Carloni et al Biorxiv 2025.04.22.649942v1) as the 177 bp repeats in *T. brucei* is untested.

**Figure 1.**
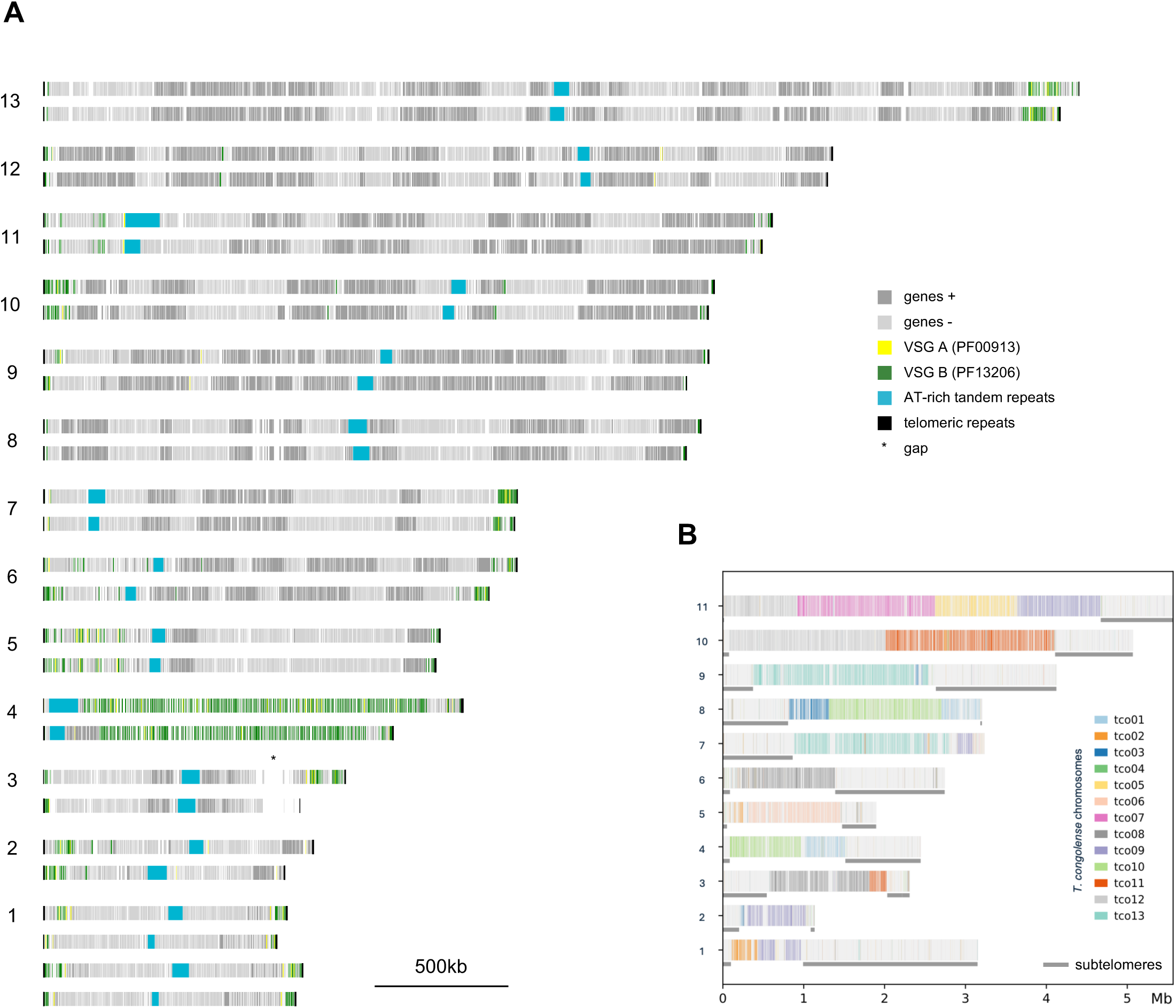
A haplotype-resolved assembly of the 13 largest chromosomes in the *T. congolense* genome and synteny with *T. brucei* megabase chromosomes. **A.** All assembled chromosomes of *T. congolense* IL3000 above 850 kbp in length are shown, comparing two haplotypes for chromosomes 2-13 and four haplotypes for chromosome 1. Genes and pseudogenes predicted using Companion^47^ are shown in two shades of grey (reflecting their coding strand), and *VSG* family genes are highlighted (yellow for Pfam PF00913, green for Pfam PF13206). An unresolved sequence gap is highlighted by an asterisk. **B.** Mapping of the newly assembled *T. congolense* chromosomes onto *T. brucei brucei* Lister 427 megabase chromosomes; *T. brucei* subtelomeres are underlined (genome version 2018 v.12)^53^.

Sequence analysis of the 28 >850 kbp contigs revealed that these represent haplotype-resolved sequences of 13 distinct chromosomes (Fig.1A, Fig.S3). 12 of the chromosomes are diploid, while one (chr 1, the smallest) is tetraploid in wild type cells. In 12 of the chromosomes directional gene clusters could be discerned (Fig.1A), consistent with the expected multigenic transcription seen in *T. brucei* and wider kinetoplastids^12,34^. In one copy of chromosome 3 a gap in the assembly could not be closed, which was centred on the repetitive splice leader gene array. In each chromosome a single locus was detected that comprised AT-rich tandem repeats, which was syntenic between the chromosome homologues but differed in size across all chromosomes (24.7-128 kbp; median 53.1 kbp, mean 55.3 kbp). It seems plausible that this feature represents a single centromere on each chromosome, consistent with what is seen in *T. brucei*^15,35,36^, but experimental evidence is needed to test such a prediction^37^.

To understand how related the large chromosomes of *T. brucei* and *T. congolense* are, we performed synteny analysis between the 11 megabase chromosomes of *T. brucei* and the 13 predicted chromosomes of *T. congolense* (Fig.1B). In no case was there complete synteny between any chromosome pairs, and instead each *T. brucei* chromosome core was a composite of at least two *T. congolense* chromosomes. Thus, there has been significant genome rearrangements since the two species emerged in evolution. In addition, there was limited evidence of sequence conservation of the *T. brucei* subtelomeres with sequences in *T. congolense* (Fig.1B), which appears consistent with previous analysis that indicated the *VSG*s of the two species are distinct in sequence and recombination^27^. Though the size range of these 13 large *T. congolense* chromosomes (0.88 -3.88 Mbp) is smaller than the megabase chromosomes of *T. brucei*, the ability to map much of the core genome content between them suggests they represent the ‘megabase’ chromosome complement of the genome. For 12/13 of the megabase chromosomes, a clear distinction from all the smaller chromosomes (see below) was the lack of discernible 369 bp repeats. The sole exception is chromosome 4, where the single AT-rich, tandemly repeated region also contains an array of 369 bp repeats. This observation may suggest that the smaller chromosomes (see below), many of which mainly contain *VSG*s, evolved from this megabase chromosome. However, the unusual composition of chromosome 4 (discussed below), may indicate that it arose by aggregation of minichromosomes and addition of a putative centromeric AT-rich locus to ensure its segregation.

### An ancestral tetraploid chromosome in *T. congolense*

Aneuploidy is common across the kinetoplastida^38^, having been documented in *Leishmania*^39,40^ and *T. cruzi*^41^, but appears scarce in *T. brucei*^42^. Much of such aneuploidy, at least in *Leishmania*, arises apparently stochastically during growth^43^, but recent work has shown that one chromosome, conserved across a wide range of trypanosomatids, is stably polyploid^44^. In *T. brucei* there no such amplified chromosome, which has instead evolved as a duplication within chromosomes 4 and 8^44–46^, thus preserving diploidy across the megabase genome. As our assembly predicted that chromosome 1 in *T. congolense* is found in four copies, we asked if this chromosome corresponds to the ancestrally duplicated chromosome by searching for orthology in *L. major* and in *T. brucei* (Fig.2A). Synteny analysis showed that the core of *T. congolense* chromosome 1 best corresponds to chromosome 31 of *L. major*, which is stably polyploid, and best synteny for this chromosome with *T. brucei* was seen on chromosomes 4 and 8, with some additional overlap with chromosome 10. Thus, *T. congolense* retains a complete chromosome that is tetraploid, suggesting an organisation more like the majority of trypanosomatids and distinct from *T. brucei*, which may therefore be unique in absorbing such raised ploidy into diploidy through duplication of the ancestral chromosome. In this regard, further analysis of the *T. vivax* genome may be warranted^44^, as well as more closely related species such as *T. simiae* and *T. suis*.

**Figure 2.**
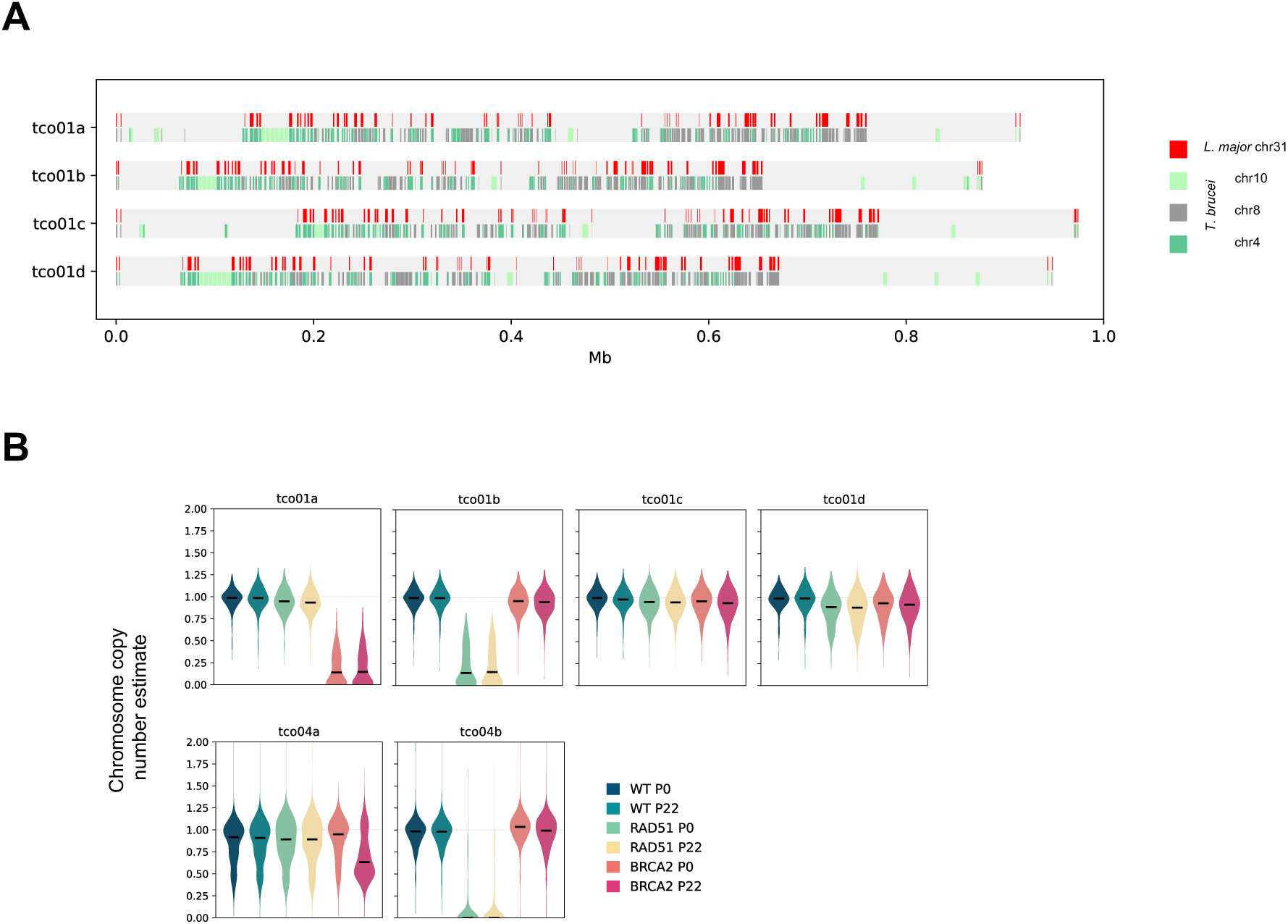
Loss of homologous recombination undermines stability of tetraploid chromosome 1 and *VSG*-rich chromosome 4 in *T. congolense*. **A.** Sequences best matching *L. major* chromosome 31, as well as *T. brucei* chromosomes 10, 8 and 4 (*T. brucei brucei* Lister 427 2018 v.12), are shown for the four copies of chromosome 1 (tco01a-tco01d). **B.** Chromosome copy number is shown for *T. congolense* IL3000 chromosomes 1 and 4 (tco01 and tco04) in wild type (WT) cells and in RAD51 or BRCA2 null mutants, before (P0) and after prolonged *in vitro* growth (P22); chromosome copy number was estimated from normalised read depth coverage of short reads across putative coding sequences.

### Loss of homologous recombination leads to ploidy reduction of select *T. congolense* chromosomes

Analysis of chromosome copy number in *T. congolense* RAD51 and BRCA2 mutants provided insight into genome stability (Fig.2B). Whereas chromosome 1 was found to be stably tetraploid in wild type cells after growth *in vitro*, a decrease in ploidy was seen in the two mutants: one copy of chromosome 1 was lost in each of the mutants. Additionally, a copy of chromosome 4 was lost in the RAD51 mutants. In contrast, no change in ploidy was seen for any other chromosome in either mutant (Fig.S4). These data suggest that functional HR is needed to maintain polyploidy of chromosome 1 for reasons that remain unclear, but it would be of interest to explore this intersection in other trypanosomatids^44^. Loss of chromosome 4 in HR mutants does not relate to polyploidy and instead may reflect *VSG* archive organisation (see below).

### Arrangement and expression of the *VSG* archive differ in *T. congolense* and *T. brucei*

To examine the organisation of *VSG* genes in the *T. congolense* genome, we localised all potential *VSG*s by identifying genes that fulfilled the following criteria: annotated by Companion^47^ as *VSG*, and contained the b-VSG Pfam domain PF13206. In total, 1529 putative *VSG*s were predicted in this way, of which Companion predicted 24.1% to be pseudogenes, comparable to that reported previously^27^ and substantially fewer than the predicted ∼80% of *T. brucei VSG*s thought to be pseudogenes^13,14^. *VSG*s were found on all megabase chromosomes (Fig.1A), and this part of the repertoire represents most (70.2%) of the *T. congolense VSG* archive. Genes containing Pfam domain PF00913 (a-VSGs in *T. brucei*) are not thought to encode VSGs in *T. congolense* but to be PAGs (procyclin-associated genes) or encode transferrin receptors^2^; these genes were less numerous than *VSG*s (Fig.5A) and were found in all the *T. congolense* megabase chromosomes (Fig.1A, Fig.5A). Given the separation of *T. congolense* VSGs into discrete phylotypes^29^, we asked if these were spatially separated across the megabase chromosomes, potentially explaining their recombinational isolation^27^, but each chromosome contained a range of phylotypes (Fig.S5B).

Arrangement of the *T. congolense VSG* archive across the megabase chromosomes was strikingly different from *T. brucei* in three ways. First, in contrast to the large *VSG*-rich subtelomeres found in each *T. brucei* megabase chromosome (median 26.7%, mean 31.1% of chromosome length)^15,16^, 12/13 *T. congolense* chromosomes contained notably smaller telomere-proximal, *VSG*-rich regions: only 0.36 to 7.55% of predicted protein-coding genes and pseudogenes in each chromosome were *VSG*s (Fig.1A, Fig.S5A). Second, the entire length of one chromosome (4) contained *VSG*s (Fig.1A): in total 533 *VSG*s were housed here, amounting to 41.2% of the total archive and representing 29% and 32% of all putative genes on the two chromosome homologues. A further measure of the abundance of *VSG*s in chromosome 4 is seen when considering only genes and pseudogenes that have identifiable Pfam domains: the VSG B Pfam domain PF13206 accounts for 347/399 and 298/331 of any detected Pfam in the two homologues (Fig.S5C). Indeed, so unusual is this organisation, and so unlike what is seen in *T. brucei*, it was impossible to derive a synteny map between *T. congolense* chr 4 and any *T. brucei* chromosome (Fig.1B). Third, analysis of sequence proximal to all 56 telomeres detected in the 28 haplotypes of the 13 chromosome homologues failed to detect any feature consistent with a *T. brucei VSG* BES: though *VSG*s were found directly adjacent to the telomere tract in 31/56 cases (see below, Fig.3), these were not preceded by a conserved set genes (based on Pfam analysis of all putative genes and pseudogenes 70 kbp upstream of telomeric repeats), and we were unable to detect 70 bp or 50 bp repeats (characteristic elements found upstream of, respectively, *T. brucei* VSGs^13,48^ and BESs^16,49^) anywhere in the *T. congolense* genome assembly. Thus, the telomeric ends of the *T. congolense* megabase chromosomes are not locations of loci resembling a *VSG* BES, which is the sole site of VSG transcription in *T. brucei* cells in the mammal^11^. Nonetheless, such sequence analysis cannot exclude that *T. congolense VSG* transcription occurs from these chromosomes in a different form of expression site, perhaps more akin to the smaller VSG expression sites used by *T. brucei* in metacyclic life cycle forms in the tsetse^50–52^.

**Figure 3.**
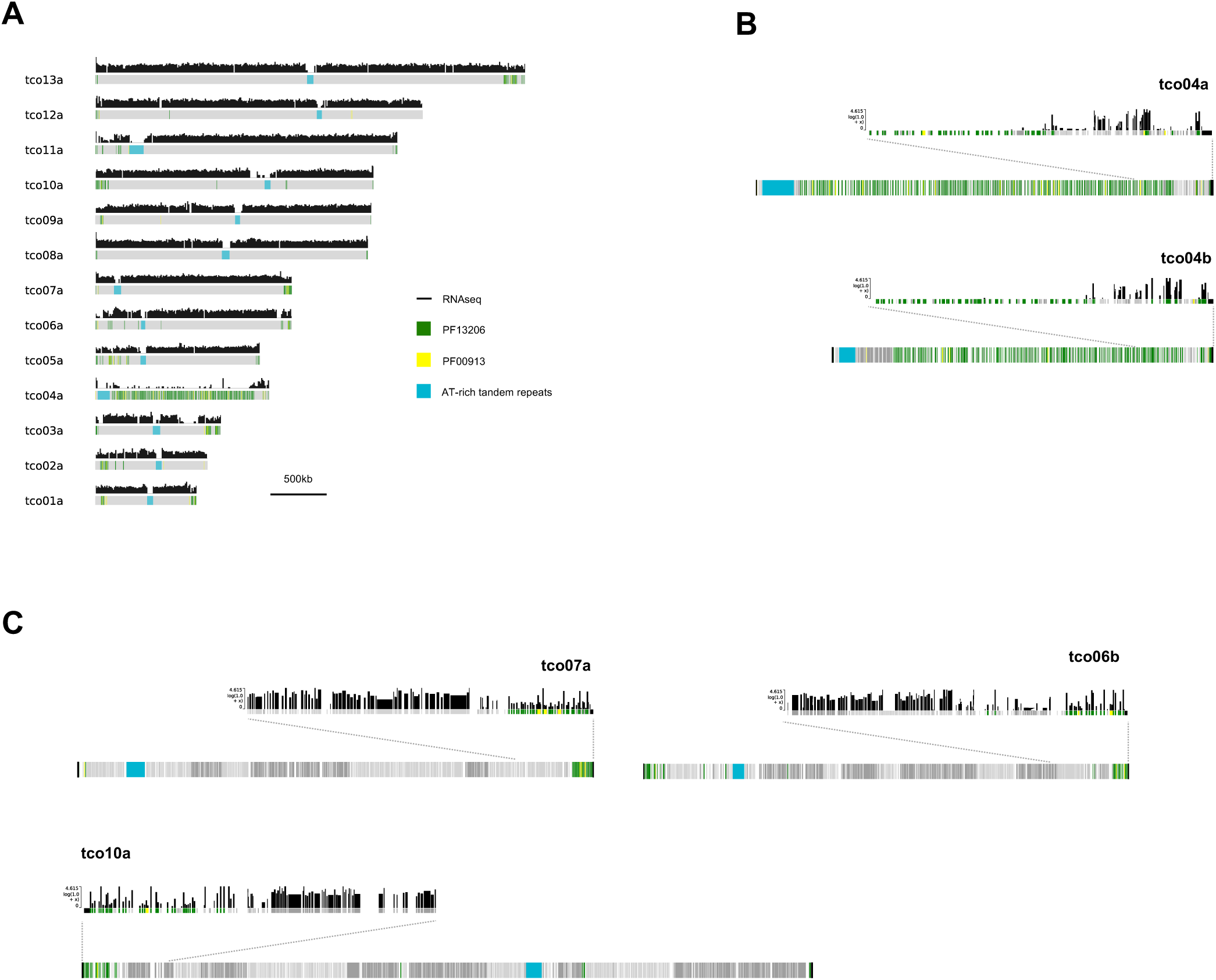
Gene expression from the 13 largest chromosomes in *T. congolense*. **A.** Bulk RNAseq data is shown mapped to haplotype A copies of the 13 largest *T. congolense* chromosomes, with VSG family Pfam-containing genes and AT-rich tandem repeat regions highlighted.; RNAseq was generated from P0 wild-type cells and analysed using Salmon (output in transcripts per million reads; TPM). **B.** A more detailed look at expression from the subtelomeric regions of both copies of chromosome 4 (tco04). **C.** As in B, a more detailed look at expression levels at subtelomeric regions of haplotype A of chromosomes 7, 6 and 10 (tco07a, tco06b and tco10a).

To understand if and how *VSG*s might be expressed in the megabase chromosomes, we mapped RNA from wild type bloodstream form *T. congolense* IL3000 to the new genome assembly: Fig.3A shows mapping to a single homologue of each chromosome, while Fig.S6 shows mapping to the two homologues of each chromosome. Consistent with the presence of multigenic transcription units, RNAseq signal was seen across most of the non-*VSG* ‘central’ regions of 12 of the 13 chromosomes; the only regions without detectable read mapping corresponded to the single putative centromeric AT-rich locus in each chromosome. Strikingly, in each of these chromosomes, RNAseq reads mapped to *VSG*s, whether distal or proximal to the telomere (Fig.3C). Applying a conservative expression threshold of minimum 5 transcripts per million (TPM), 309/540 (57.2%) of the putative *VSG*s on 12/13 chromosomes were considered expressed. Chromosome 4 was again a notable variant: here, there was little evidence for RNAseq mapping across most of the chromosome (Fig.3A,B, Fig.S6) and, applying the same threshold, only 21/533 (3.94%) *VSG*s were expressed, with the clearest RNAseq signal being telomere-proximal. Whether this single chromosome acts as a silent *VSG* reservoir comparable to that spread across all subtelomeres of *T. brucei* and thereby provides a source of new VSG variants by recombination, remains to be seen. Equally, what purpose(s) is served by the many other unexpressed non-*VSG* genes in chromosome 4 is unclear. Nonetheless, taken together, these data provide no evidence for predominant transcription of *VSG* from a single locus in the megabase chromosomes, though it remains possible that telomere-proximity favours VSG expression.

### Hi-C provides no evidence for distinct core and subtelomere genome compartments in *T. congolense*

Assembly of the *T. congolense* genome was informed by Hi-C interaction data, and so we next analysed these data in more detail to ask if the genome of *T. congolense* is organised within the nucleus like that of *T. brucei*. Fig.4A shows Hi-C interaction matrices of inter- and intra-chromosomal DNA interactions for all megabase chromosomes; these data support the assembly and arrangement of these larger chromosomes. To investigate whether large-scale genome compartmentalisation can be seen in *T. congolense*, as described in *T. brucei*^15,53^ and *T. cruzi*^54,55^, we generated correlation matrices, along with 1-10 eigenvector data and insulation scores, and measured contact decay, from the Hi-C data (Fig.4B, Fig.S7A,B). Distinct genomic compartments could not be clearly discerned in *T. congolense* from any of these analyses, an observation that aligns with the above RNAseq data (Fig.3, Fig.S6), which failed to reveal a clear difference in transcription between the subtelomeric, *VSG*-rich parts of the chromosomes and in the *VSG*-denuded chromosome cores.

**Figure 4.**
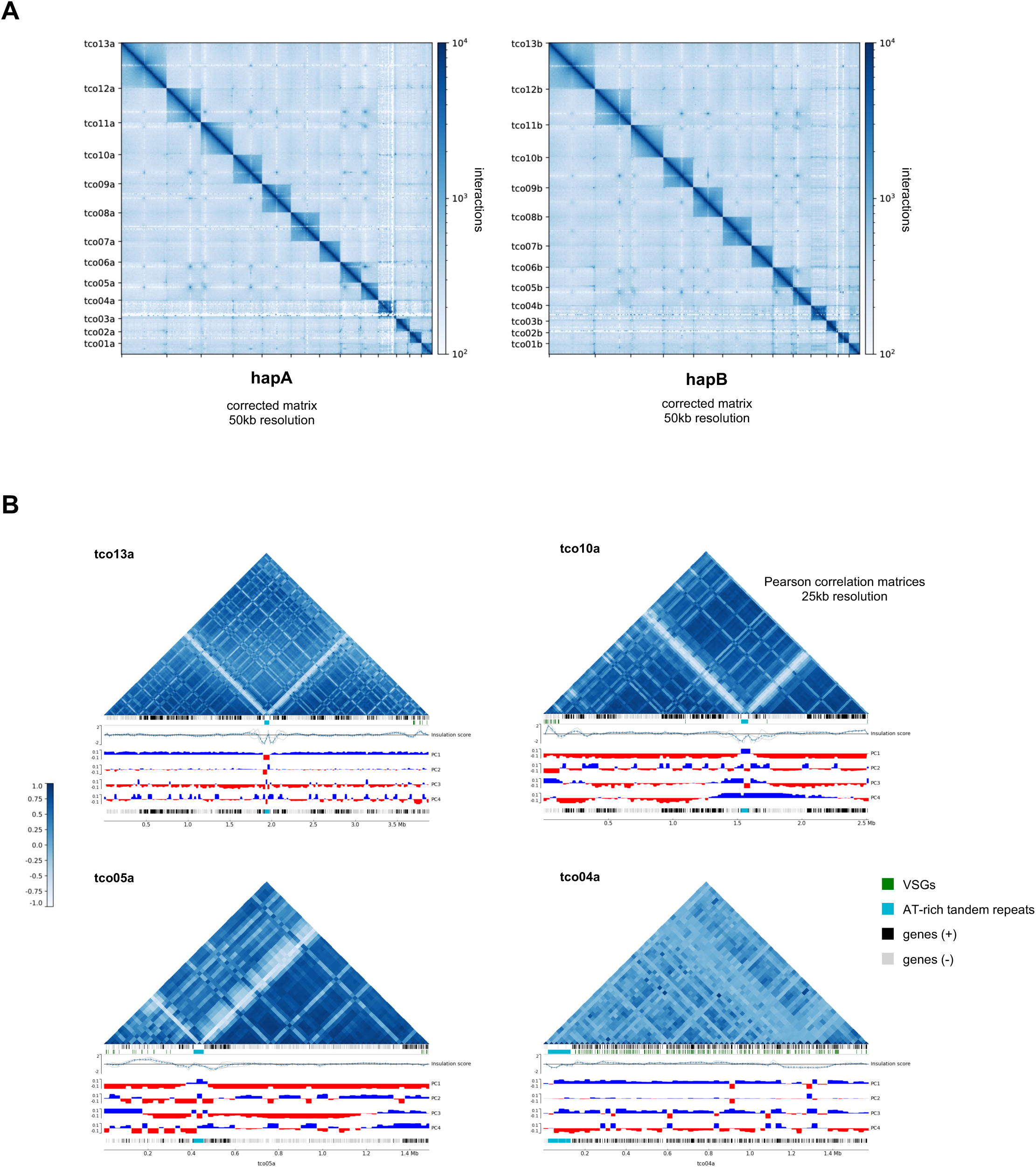
Hi-C provides no evidence for large-scale compartmentalisation within the 13 largest *T. congolense* chromosomes. **A.** Corrected Hi-C matrix for haplotype A and haplotype B of the 13 largest *T. congolense* chromosomes at 50 kbp resolution (interaction matrices not corrected for distance decay). **B**. Examples of correlation matrices, along with insulation scores and 4 eigenvector values, for four chromosomes at 25 kbp resolution. Putative genes are highlighted in black or grey, depending on coding strand; *VSG* genes are highlighted in green and putative centromeres in light blue.

**Figure 5.**
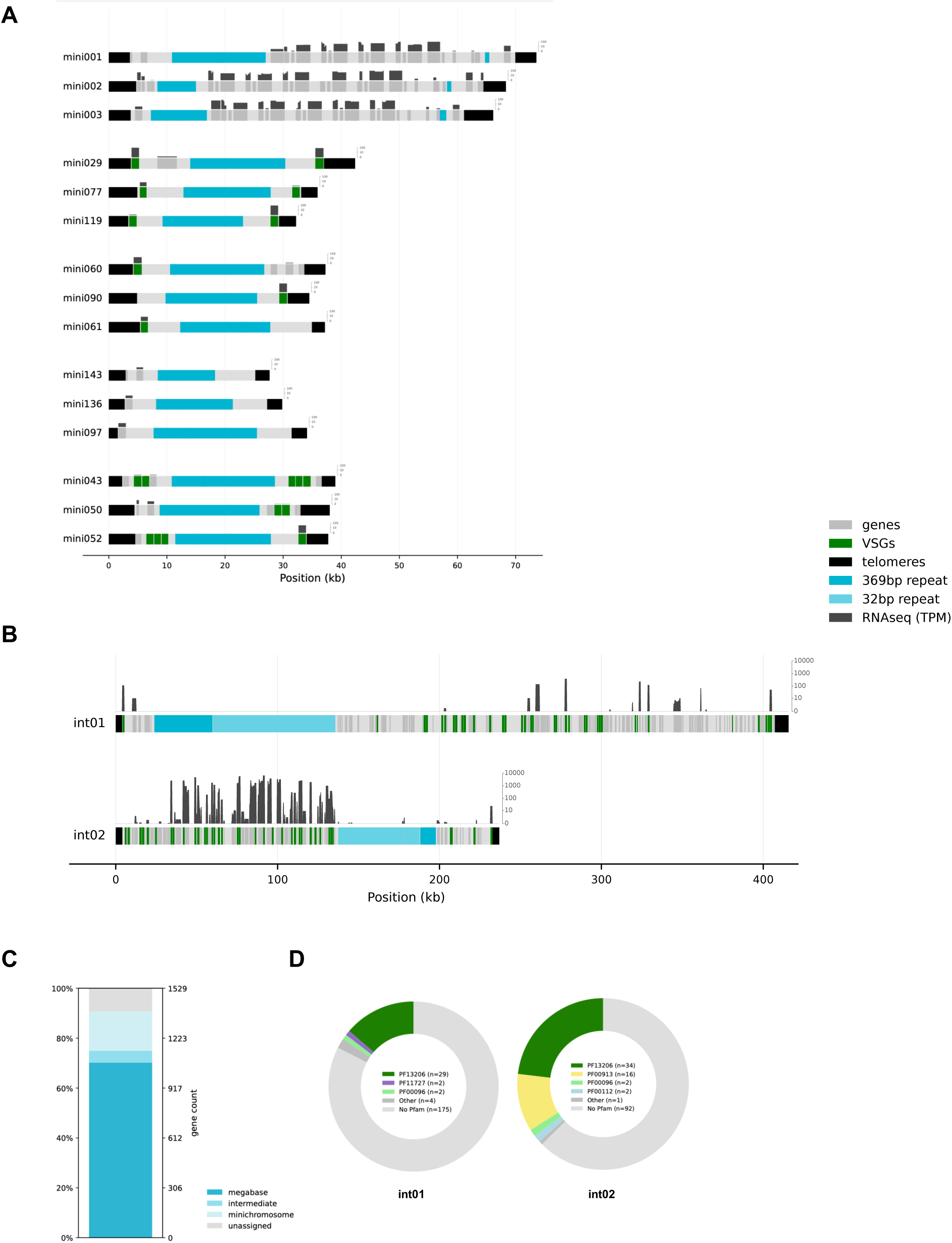
Assembly and expression of the abundant small chromosomes in the *T. congolense* genome. **A.** Examples of *T. congolense* IL3000 minichromosomes and their gene expression from bulk RNAseq data in wild type cells, with putative genes, *VSG*s and repeat elements highlighted. **B.** as in A but for two predicted intermediate chromosomes. **C.** The distribution of putative *VSG* genes across the different chromosome groups (megabase: chromosomes >850 kbp; intermediate: 200-500 kbp; and minichromosomes: <100 kbp). The genome assembly also contains contigs that are not telomere-to-telomere assemblies, and these are categorised as ‘unassigned’. **D.** Gene content of the two intermediate chromosomes shown in B, based on Pfam analysis of translated putative gene and pseudogene sequences.

Two *T. congolense* genomic elements did, however, stand out in the Hi-C analyses: the AT-rich locus on each chromosome, and sites of gene direction change (Fig.4, Fig.S7B, Fig.S8). In *T. brucei*, centromeres have been shown to act as pronounced boundaries within the megabase chromosomes^15^, and the interaction profile around the AT-rich locus on each *T. congolense* chromosome (Fig.4B, Fig.S7B) may be consistent with these being centromeres. However, we note that high conservation of the AT-rich repeats in these loci may confound the analysis. While no transcription initiation- or termination-associated factors, including histone variants, have been mapped to the *T. congolense* genome, many of the loci inferred to be the ends of directional gene clusters, by virtue of change in common gene orientation, showed shared interaction patterns (Fig.4B, Fig.S7B). This observation shares a similarly with what has been observed at transcription start sites in *T. brucei*^53^, suggesting transcriptional hubs are a broad feature of trypanosomatid nuclei, driven by the needs of widespread polycistronic transcription. Notably, such interaction profiles were less clear across chromosome 4 (Fig.4B), consistent with RNAseq evidence for a lack of gene expression in the *VSG*-rich core.

### Organisation of and VSG expression from the abundant small chromosomes in *T. congolense*

The above discussion focuses on the megabase chromosomes of *T. congolense*, but the assembly additionally predicted 147 telomere-telomere chromosomes ranging in size from 27-416 kbp (Fig.5). Based on their gene content, size, chromosome organisation and repeat content, these smaller chromosomes can be split in three groups. In the first group, consistent with previous reports^23^, the 141 smallest chromosomes (27-55 kbp) contain a central core of tandemly repeated 369 bp repeats, flanked by one or more putative coding sequences; 135/141 of these chromosomes contain 1-5 putative VSGs (Fig.5A). The second group, comprising four slightly larger (64-74 kbp) and less symmetric chromosomes, each has a large and small gene coding region (40-48 kbp) flanking the 369 bp repeat region (Fig.5B), and no detected *VSG*s. The third group comprised two notably larger chromosomes (237 and 416 kbp) that contain an additional AT-rich region adjacent to the 369 bp repeat region (Fig.5C) and, based on Pfam content analysis, contain predominantly genes with no detected Pfam domains, as well as numerous VSG A and VSG B family genes (Fig.5D). This variable composition of submegabase chromosomes in *T. congolense* warrants comparison with a recent re-evaluation of the small chromosomes of *T. brucei*^16,20^, in that in both species they are not solely repositories of *VSG*s but contain other genes and, furthermore, their evolution has involved retention of likely centromeric elements to allow DNA replication and/or provide mitotic stability.

In *T. brucei*, no work has documented *VSG* expression from minichromosomes^56^, whereas a previous report suggested sequence elements in some *T. congolense* submegabase chromosomes that might allow telomere-adjacent transcription^23^. To test this proposed difference in function, Fig.5A,B shows RNAseq mapping to the submegabase chromosomes assembled here. These data reveal that *T. congolense* small chromosomes display *VSG* expression: using the same TPM cut-off of minimum 5 TPM, 121/239 (50.63%) putative *VSG*s located on these chromosomes are expressed across 97/135 *VSG*-containing small chromosomes. Furthermore, at least in the two largest small chromosomes, transcripts were detected from non-telomeric *VSG*s (Fig.5B). Thus, this component of the *T. congolense* genome reveals further the disparity between *T. brucei* and *T. congolense* VSG expression dynamics.

### Conclusions

A combination of long-read ONT sequencing and Hi-C DNA interaction analysis has revealed the genome of *T. congolense* to be composed of 12 diploid chromosomes, one tetraploid chromosome, and more than 100 small chromosomes. Superficially, this organisation compares with the similarly sized genome of *T. brucei* and indicates that both genomes have been shaped by the need for antigenic variation, in that *VSG*s are the main gene found on most small chromosomes, and a large number of *VSG*s localise to the subtelomeres of most of the larger chromosomes. However, a deeper analysis reveals several differences, which indicate deviation in the way *VSG* expression is organised in the two parasites: telomere-telomere assembly of all chromosomes confirms previous suggestions^23^ that *T. congolense* does not contain *T. brucei*-like VSG expression sites; *VSG*-rich subtelomeric regions of the largest chromosomes in *T. congolense* are notably smaller than seen in *T. brucei*; *VSG* expression in *T. congolense* is not limited to telomere-adjacent genes, consistent with a lack of subnuclear compartmentalisation between chromosome cores and subtelomeres; a single *T. congolense* chromosome acts as a potential silent reservoir of *VSG*s; and *VSG* expression can be detected from multiple small chromosomes. Thus, this genome assembly provides a comprehensive platform to dissect the operation of host immune evasion by VSG expression in *T. congolense*.

## Methods

### Parasite culture, generation of mutant lines

Bloodstream form *T. congolense* strain IL3000^57^ was used for all experiments (originally isolated at the International Livestock Research Institute (Nairobi, Kenya) and derived from infected mouse blood, received from Theo Baltz, University of Bordeaux). Parasites were cultured at 34 °C with 5% CO_2_ in modified HMI-93^58,59^ media, with 10% goat serum (Gibco) and 20 g/L albumax II (Thermo-Fisher). Cells were routinely cultivated in 6 or 24 well plates and a P1000 pipette tip or 5ml stripette, as appropriate, was used to flush the cells adhered to the bottom of the well prior to passage into a new well, or for haemocytometer counting of cells. For large volumes, 75 cm^2^ flasks were used, and a 10 ml or 25 ml stripette used to flush the parasites sticking to the surface of the flask.

To generate *RAD51* (TcIL3000.11.8740) and *BRCA2* (TcIL3000_1_180) null mutant parasites, plasmids pJM-RMC-09 (TcoRAD51) and pJM-RMC-10 (TcoBRCA2) were synthesised by swapping the UTRs of *RAD51* and *BRCA2* into pJM-RMC-03 (BSD-resistance; Addgene plasmid # 248661; https://www.addgene.org/248661/), using restriction enzyme cloning. Briefly, UTRs were PCR-amplified from *T. congolense* IL3000 gDNA with Phusion polymerase (NEB UK, M0530S; see Table S2 for all primer sequences) and ligated into pGEM-T Easy Vector (Promega UK, A1360). Correct amplification of the UTRs was confirmed by Sanger Sequencing (Eurofins Germany) with primer M13F. To insert the 5’ UTRs, pJM-RMC-03 and both pGEM-T Easy with 5’ UTR plasmids were digested with *Not*I-HF and *Hind*III-HF (NEB UK, R3189S and R3104S) and ligated with T4 ligase (Promega UK, M1801). To insert the 3’ UTRs, the resulting plasmids and pGEM-T Easy 3’ UTR plasmids were digested with *Nsi*I-HF and *Xho*I (NEB UK, R3127S and R0146S) and ligated together. Correct insertions were confirmed by Sanger Sequencing (Eurofins Germany; 5’UTR confirmation: JM015; 3’UTR confirmation: SP6). The hygromycin-resistance plasmids pJM-RMC-19 (RAD51) and pJM-RMC-20 (BRCA2) were generated by restriction enzyme cloning; briefly, plasmids pJM-RMC-09 and pJM-RMC-10 and pyrFEKO-HYG (a gift from George Cross; Addgene plasmid # 24020; https://www.addgene.org/24020/) were digested with *Bgl*II and *Xba*I (NEB UK, R0144S and R0145S), with the backbones minus antibiotic ligated to the hygromycin resistance gene from pyrFEKO-HYG using T4 ligase. Correct integration of the new antibiotic resistance gene was checked using Sanger Sequencing (primer JM042).

To generate the null mutant parasites, the relevant plasmids (pJM-RMC-09, -10, -19 or -20) were linearised using *Not*I-HF and *Xho*I (NEB UK). Around 3-4 ×10^7^ *T. congolense* cells were flushed, counted and spun at 1250 *g* for 10 minutes. They were then resuspended in 100 µl Tb transfection buffer^60^, mixed with 10 µg of the relevant digested DNA and electroporated in an Amaxa Nucleofector II (Lonza) using program Z-001. Parasites were resuspended in 20 ml of fresh media and immediately plated out at 1:5, 1:10 or 1:20 dilutions n 24 well plates, and with the remaining pooled cells plated in 6 well plates. After around 18 hours, antibiotic selection was commenced, with final concentrations of 0.25 μg/ml hygromycin B (Roche, 10843555001) or 0.5 μg/ml blasticidin (Invivogen, ant-bl-1), depending on the linearised plasmid used. Successful integration of the antibiotic-resistance cassettes was assessed by PCR across the relevant locus: gDNA was extracted using a Qiagen DNeasy Blood and Tissue Kit, and Phusion polymerase used to amplify target fragments, Fig.S1). RT-qPCR of the double antibiotic-resistant clones confirmed RNA loss: RNA was extracted using a Qiagen RNeasy Kit, cDNA produced using Superscript IV (Thermo Fisher), and an Applied Biosystems 7500 Fast machine used with Fast Sybr Green master mix (Thermo Fisher), all according to the manufacturer’s instructions. The lines generated were designated TcoRAD51^−/-^clone H2B2 and TcoBRCA2^−/-^ clone H4B1.The TcoBRCA2^−/-^ H4B1 line had the antibiotic resistance cassettes removed by the transient expression of pLEW100cre (a gift from John Donelson via George Cross; tryps.rockefeller.edu/Plasmids/pLEW100cre.txt) and induction of Cre-recombinase using 1 µg/ml tetracycline (Merck, T7660) as per Scahill et al^61^, to generate TcoBRCA2^−/-^ floxed clone 2 (fx2). Parasites were cloned in a 24-well plate (at 1:10 dilution, straight after transfection) and selected using 100 µg/mL ganciclovir (Merck, G2536) 16 hours post transfection. Clones were checked for sensitivity to blasticidin and hygromycin.

### Cloning of Parasites

*T. congolense* IL3000 WT, TcoRAD51^−/-^ clone H2B2 and TcoBRCA2^−/-^ fx2 parasites were recloned, by dilution of each line to 0.5 cells/well in a single 24 well plate. Plates were monitored for growth and at least three clones per line were selected for growth. The starting clones (P0) were stabilated as soon as possible after cloning (once a density of ∼1-2 × 10^6^/ml had been achieved in a well of a 6 well plate; around 22-28 generations from a single cell; depending on the line). Stabilates were made using a final concentration of 15% glycerol in HMI-93 media. The three clones were then sub-passaged 2-3 times per week, growing in 6 or 12 well plates to a density around 2-3 × 10^6^/ml,before being diluted to around 5 × 10^4^/ml at each passage. After 11 passages, stabilates were made of each clone (P11 samples; ∼80-90 further generations from the starting culture), followed by a further 11 passages, and stabilates made of the surviving clones (P22 samples; ∼80-90 generations from the P11 samples). Three clones for each line survived to passage 22; two per line were used for further analysis (WT IL3000 clones 1 and 2; TcoRAD51^−/-^ clones 2 and 5; TcoBRCA2^−/-^ clones 1 and 2).

### Nucleic Acid extraction and sequencing

Stabilates of each clone at each passage time point were removed from liquid nitrogen and recovered in 6 ml media in one well of a 6 well plate. Once growing, these parasites were transferred to four 75 cm^2^ filter top culture flasks, with 40 ml HMI-93 media and grown on their back to allow attachment of the parasites on the back surface. Parasites were grown to a density of between 2 and 5 × 10^6^/ml, before processing. For RNA extraction, 6×10^7^ cells were pelleted from each individual flask (giving 4 replicates), and RNA extracted from each pellet using a Qiagen RNeasy Extraction kit, following manufacturer’s instructions, and including the on-column DNase-digestion steps. RNA libraries were prepared from the three samples with the highest RNA concentration using an Illumina TruSeq stranded mRNA prep kit and an Illumina NextSeq2000 was used to perform paired-end sequencing (2×100bp). The libraries were sequenced by the Molecular Analysis facility at MVLS Shared Research Facilities (Glasgow University).

For ONT sequencing, high molecular weight genomic DNA was extracted from each of the six clones at both P0 and P22 using the Monarch HMW DNA Extraction Kit for cells and blood (NEB UK), pooled from the same flasks as used for the RNA extractions, with lower density flasks refilled with media and grown for a further 2-3 days. Around 1.5 – 2 ×10^8^ cells were pelleted by centrifugation, washed with PBS, and stored at -80 °C before DNA preparation. During library preparation, ONT Ligation Sequencing Kit (SQK-LSK114) was used as per manufacturer’s instructions, and sequencing was performed on a GridION Mk1 device using R10.4.1 MinION flow cells. The same high molecular weight gDNA was also submitted to BGI Genomics for short read sequencing using DNBseq paired-end sequencing (2×100bp), with 30M reads per sample. Six samples were sequenced: IL3000 WT clone 2 P0 and P22; TcoRAD51^−/-^ clone 5 P0 and P22, and TcoBRCA2^−/-^ clone 1 P0 and P22. For PacBio sequencing, six C57BL/6J mice were infected with WT IL3000. At day 7 post infection, all six were culled and their blood combined. The parasites were purified through a DE52 column^62^, washed with PBS-G and pelleted by centrifugation at 600 *g* for 10 mins, washed in PSG buffer, pelleted again and stored at -80 °C. DNA was extracted from thawed pellet using the Monarch HMW DNA Extraction Kit for cells and blood kit, using the manufacturer’s instructions. High molecular weight DNA was submitted to the Earlham Institute for library preparation (using Express Template Prep Kit), and sequencing using PacBio Sequel II SMRT cell.

For Hi-C, IL3000 WT clone 1 was used, five further passages from the starting clone P0 culture described above (around 40 generations). The Proximo Hi-C (Microbe) Kit from Phase Genomics (Seattle, USA) was used, following the manufacturer’s instructions. Two samples were prepared, from 6.5 × 10^8^ cells and 5 × 10^8^ cells respectively. The samples were combined after the DNA purification steps of the protocol and one library made using both. An Illumina NextSeq2000 was used to perform paired-end sequencing (2×100 bp), by the Molecular Analysis facility at MVLS Shared Research Facilities (Glasgow University).

### ONT data pre-processing

ONT whole genome sequencing data was basecalled using dorado (v.0.9.1, https://github.com/nanoporetech/dorado) duplex sup and samtools (v.1.19.2)^63^ was used to split the output reads into simplex, duplex and de-duplexed simplex reads for further processing based on the dx tag value. Chopper (v.0.9.0)^64^ was used to remove ONT control sequence (DCS) reads. Dorado correct was used to correct the simplex data, which was used for *de novo* genome assembly downstream.

### Genome assembly and QC

A combination of two *de novo* genome assemblies of *T. congolense* IL3000 bloodstream-form cells was used to create the genome version presented here. Verkko (v.2.2.1)^32^was used to assemble the haplotype-resolved megabase chromosomes of the parasite and hifiasm (v.0.25.0)^65^*de novo* assembly was used for the smaller chromosomes, as these were not assembled by verkko. For both assemblies, the same sets of data were used: Hi-C reads, dorado-corrected ONT simplex data (provided as ‘--hifi’ in verkko) and uncorrected ONT simplex data (as ‘--nano’ in verkko and ‘--ul’ in hifiasm). A number of other genome assembly approaches were tested with the available data as well (Fig.S2).

For quality control (QC), a number of metrics were used to evaluate the generated genome assemblies: BUSCO (v.5.8.3)^66^ was used to determine and compare genome completeness (euglenozoa_odb10 database used); consensus quality (QV) was assessed using merqury (v.1.3)^30^ and meryl (v.1.4.1) (k-mer size 17), and QUAST (v. 5.3.0)^67^ was used to evaluate general genome metrics (gene detection using glimmer, rRNA detection enabled). Telomere-to-telomere (T2T) assessment was carried out by finding telomeric (’TTAGGG’’) repeats using seqtk telo (v.1.4) ( https://github.com/lh3/seqtk) and homologous chromosomes (both within an assembly and between assemblies) were identified using nucmer and promer (mummer suite, v. 3.23)^68^ with the ‘–maxmatch’ flag. The same approach (promer with ‘--maxmatch’) was used, reciprocally, to find synteny with *T. brucei* and *L. major* genomes.

### Genome annotation

Putative genes and pseudogenes were annotated using Companion (https://companion.gla.ac.uk/)^47^ with the following settings: reference protein alignment enabled, RATT conserved gene transfer on species level enabled, taxon ID 5654. Repetitive genome elements were identified using Tandem repeats finder (TRF) (v.4.09.). Repeat motifs (369 bp, 32 bp, 70 bp and 50 bp) were located or searched using fimo of meme suite of tools (v. 5.5.9)^69^ with the p-value cutoff of 10-^9^.

Putative variant surface glycoprotein (VSG) family genes were identified based on the presence of the Pfam domains and companion gene annotation: *VSG* genes, Pfam domain PF13206; other VSG family genes, Pfam domain PF00913 (identified in translated predicted gene and pseudogene sequences using hmmsearch (v.3.4) (hmmer.org), and only significant hits were retained. For this, gene and pseudogene DNA sequences were translated using bedtools getfasta (v.2.31.1)^70^ and seqkit translate (v.2.8.2)^71^. To further profile *VSG* diversity, hmm profiles of described *VSG* phylotypes^72^ were used to detect phylotype-specific protein signatures in the translated gene and pseudogene sequences (hmmsearch was used, as above, retaining only significant hits). Pfam domain searches across the translated genes and pseudogenes were performed using pfam_scan (https://github.com/aziele/pfam_scan), utilising InterPro Pfam domain motif data (https://www.ebi.ac.uk/interpro/download/pfam/) and Pfam-defined cutoffs for the individual motifs. Genome-wide sequence composition was assessed using bedtools makewindows and bedtools nuc (v.)^70^ in 1 kbp windows; this was also used to identify AT-rich repetitive regions found on the larger chromosomes.

### Chromosome copy number analysis

Chromosome copy number was estimated utilising short read whole genome sequencing data similarly to previously published work^73^ Briefly, paired-end reads were mapped to the genome assembly using bwa mem (v.0.7.19)(https://arxiv.org/abs/1303.3997) and samtools (v.1.22) was used to sort, index and remove duplicate reads from the resulting alignment files. BamCoverage from the deepTools package (v.3.5.5)^74^ was used to calculate read depth across the genome (normalisation using RPKM, 50 bp bins, reads centred). The resulting bedgraph files were used in a custom python script that extracts coverage across all annotated genes and pseudogenes and calculates the mean coverage per gene; this was then normalised against the median coverage of all genes in the genome. Per-chromosome copy number was computed as the median of these normalised values for a given chromosome.

### RNAseq analysis

RNAseq sequencing data was trimmed using Trim Galore! (v.0.6.10, paired end settings, fastQC output enabled) (https://github.com/FelixKrueger/TrimGalore) and salmon quant (v.1.10.3., automatic library type detection enabled)^75^was used to quantify the RNAseq data in transcripts per million (TPM); for running salmon quant, salmon index was run in decoy-aware mode as recommended by its developers, by running the generateDecoyTranscriptome.sh script. Both genes and pseudogenes, as predicted using Companion annotation, were used as the transcriptome.

### Hi-C data analysis

HiCExplorer (v.3.7.6)^76^ was used to analyse the Hi-C data, as per the developers’ recommendations (https://hicexplorer.readthedocs.io/en/latest/content/example_usage.html). Briefly, the genome was split into two haplotypes (A and B), which were analysed separately; bwa mem with the recommended settings was used to map the reads (-A 1 -B 4 -E 50 -L 0) and samtools was used to change the output file format to bam. Following this, hicBuildMatrix was used to generate matrices at 5 kbp resolution, and was subsequently merged using hicMergeMatrixBins to generate 10 kbp, 25 kbp, 50 kbp and 100 kbp resolution matrices. HicCorrectMatrix was used to remove outlier values (ICE, thresholds used –2.0, 5.0). HiCExplorer hicPCA was used to generate Pearson and Obs/Exp matrices, as well as retrieve eigenvectors 1-10 across the 5 resolution corrected matrices.

Topologically associated domains (TADs) and genome-wide insulation score were called using hicFindTADs (FDR correction, --minDepth 3 bins, --maxDepth 10 bins) and hicPlotDistVsCounts was used to investigate the distribution of contacts with distance between the two haplotypes at 25 kbp, 50 kbp and 100 kbp resolutions.

## Supporting information

All supplementary Figures and a Supp Table

## Data availability

Sequencing reads will be deposited to the NCBI Sequence Read Archive (SRA), and the assembled genome will be available at the EMBO-EBI European nucleotide archive (ENA).

## Author contributions

MK, JM, SL, LM, KM, RM wrote and edited the manuscript. MK, JM, DB, GO, SL, CL were responsible for methodology, investigation and analysis, including software. LM, KM, RM, supervised and conceptualised the project, and secured funding.

## The authors declare no competing interests

## Acknowledgments

We thank Zoha Khan and Kehinde Foluke Paul-Odeniran for assistance with producing *T. congolense* plasmids, and the Molecular Analysis Facility of the MVLS Shared Research Facilities for sequencing assistance. This work was supported by the Wellcome Trust (224501/Z/21/Z to RM, 221717/Z/20/Z to KM, 206815/Z/17/Z to KM, LM and RM), and the BBSRC (BB/W001101/1 to RM); the Roslin Institute (LM) is supported through core funding from the BBSRC (BS/E/D/20002173; BBS/E/RL/230002C) .

## Supplementary figure and table legends

**Figure S1. Confirmation of TcoRAD51 and TcoBRCA2 null mutants. A.** Diagram illustrating the insertion of pJM-RMC-09 or -19 into the native *TcoRAD51* locus; primer binding sites are indicated by coloured arrowheads, and the gel image shows PCR using these primers to test for loss of the TcoRAD51 ORF in wild type cells (WT), single allele knockout cells (sKO), and double allele knockout cells (dKO). **B.** As in A, but for *TcoBRCA2*. **C.** RT-qPCR of *TcoRAD51* and *TcoBRCA2*, showing expression of each gene relative to that in WT for both TcoRAD51^−/-^ H2/B2 and TcoBRCA2^−/-^ Fx 2 (double allele KO cells). The mean of 4 experiments (each of 3 replicates) is shown ±SD.

**Figure S2. Summary of evaluated genome assemblies.** During the genome assembly process, several assemblies were produced using a range of data combinations, assembly tool settings, and two assemblers. Numbers 1-15 refer to the combinations used. **A.** The upset plot highlights the various combinations of data and settings used in assembly attempts for *T. congolense*, showing the two assemblers – hifiasm and verkko – in separate panels. P0 refers to refers to long read ONT data collected from wild type cells at passage 0 (prior to prolonged *in vitro* growth of wild-type clones 1 and 2). **B.** Breakdown of how many contigs in each size category were retrieved in each assembly attempt, and whether the contigs contained both, one, or no telomeric ends (TTAGGG), based on the output of seqtk telo. The y axis represents the different assemblies, 1-15. **C.** BUSCO results. **D.** QV scores for the 15 assemblies.

**Figure S3. Chromosome pairing of the largest 13 *T. congolense* chromosomes.** The figure highlights the best within-genome matches for the largest 28 contigs (excluding self-matches), allowing the identification of homologous chromosome pairs (data generated using nucmer of mummer suite of tools, and identical top matches in 1 kbp bins have been merged). The four copies of chromosome 1 (tco01a-tco01d) have an unusual appearance due to different regions of the contigs having different best matches among the other three copies. Chromosome 4 (Tco04a and tco04b) shows more limited homology compared to other chromosomes.

**Figure S4. Chromosome copy analysis for chromosomes 2-3 and 5-13.** Panels show the estimated chromosome copy number for chromosomes not shown in Figure 2. Data for wild-type, *BRCA2*-null and *RAD51*-null cells is shown; P0 and P22 refer to passages during prolonged *in vitro* growth.

**Figure S5. *VSG* family genes in the largest 13 *T. congolense* chromosomes. A.** Distribution of VSG family Pfam-containing genes and pseudogenes in the larger chromosomes (PF00913, VSG A; PF13206, VSG B Pfam domains). **B.** Distribution of phylotypes among the putative *VSG* genes (based on VAPPER Pfam domains^72^). The ‘undetermined’ category contains sequences that either do not have a significant phylotype motif hit, or more than one motif is detected above the significance threshold. The number of *VSG*s for each chromosome contig is shown above the plot. **C.** Gene content of the two haplotyps of chromosome 4 (tco04a and tco04b) based on Pfam analysis; numbers in brackets indicate the number of genes and pseudogenes, as determined by pfam_scan (note – pfam_scan uses different thresholds for detecting sequence motifs than hmmer, which was used for detecting Pfam domains for panels A and B).

**Figure S6. Gene expression of the largest *T. congolense* chromosomes.** Bulk RNAseq data is shown mapped to all copies of the largest *T. congolense* contigs (>850 kbp), with VSG family Pfam-containing genes and AT-rich tandem repeat regions highlighted. RNAseq was obtained from wild-type cells and analysed using Salmon (output in TPM).

**Figure S7. Contact decay between chromosome homologues and correlation matrices for chromosomes 1-3, 6-9, 11-12. A.** Contact decay for haplotype A (hapA) and haplotype B (hapB) chromosome contigs at different resolutions (25, 50 and 100 kbp, as plotted by hicPlotDistVsCounts of HiCExplorer suite of tools). **B.** Correlation matrices at 25 kbp resolution for haplotype A chromosome contigs not presented in Figure 4B.

**Figure S8. Corrected interaction matrices for the 13 largest chromosomes of *T. congolense*.** Corrected matrices are shown at 25 kbp resolution for all haplotype A chromosome contigs (tco01a-tco13a), with predicted genes highlighted in black and grey (gene strand), putative *VSG*s shown in green, and AT-rich tandem repeats (putative centromeric repeats) shown in light blue. Data filtered out due to multimapping appears in white.

**Table S1. QUAST metrics for *T. congolense* IL3000 genome assemblies.** Various genome assembly metrics, as reported by QUAST, are shown for the two previously published *T. congolense* IL3000 genomes (‘IL3000’ and ‘IL3000_2019’), as well as the genome assembly presented here (‘2026’). ‘IL3000’ corresponds to TriTrypDB build version 68 assembly ‘TriTrypDB-68_TcongolenseIL3000’, and ‘IL3000_2019’ corresponds to ‘TriTrypDB-68_TcongolenseIL3000_2019’.

**Table S2.**
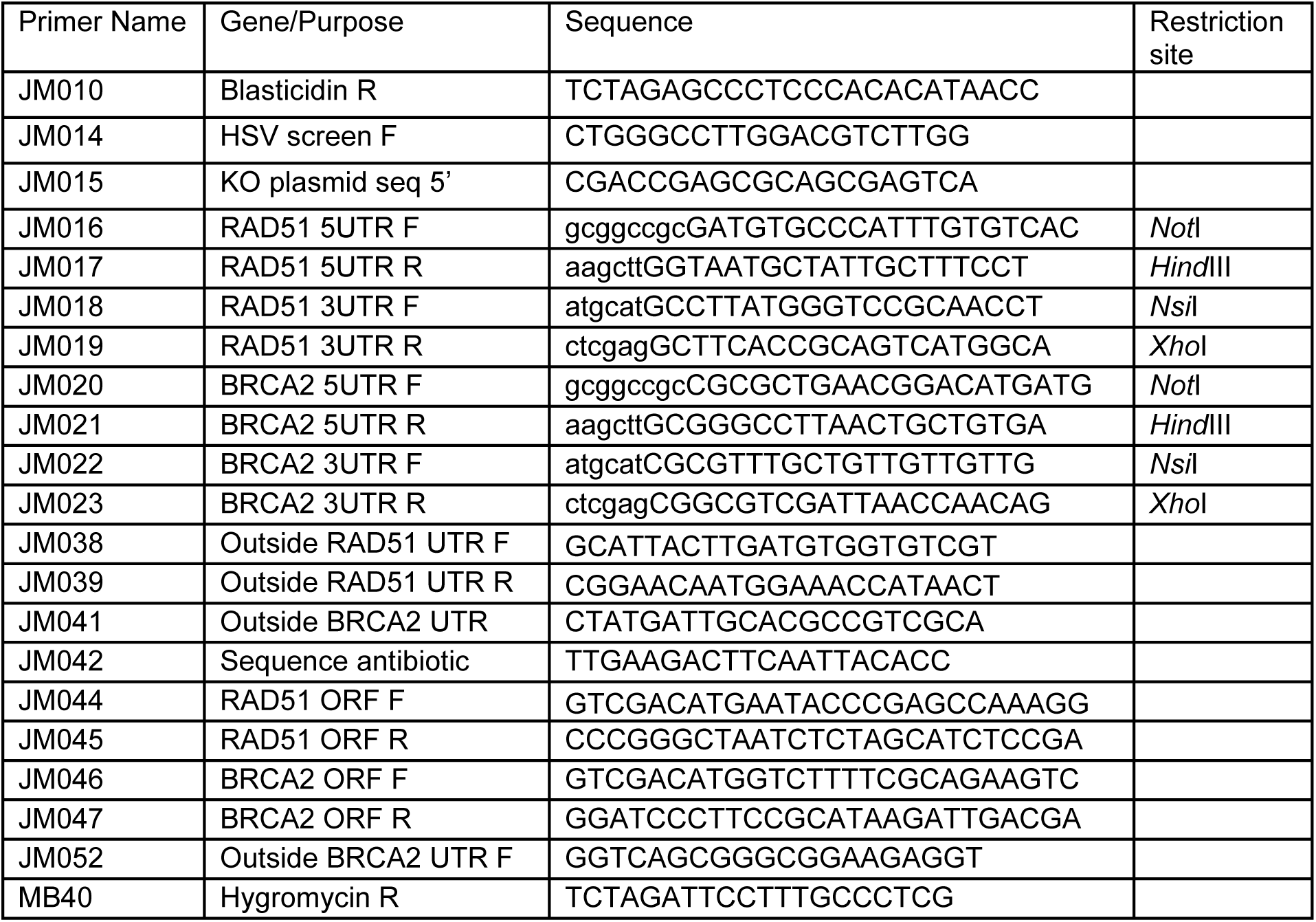
Primers Used.

