## Supplementary material for "De novo assembly of the *Trypanosoma congolense* genome reveals an organisation influenced by antigenic variation but distinct from *Trypanosoma brucei*": All supplementary Figures and a Supp Table

### Figure S1

**A** TcoRAD51$^{-/-}$

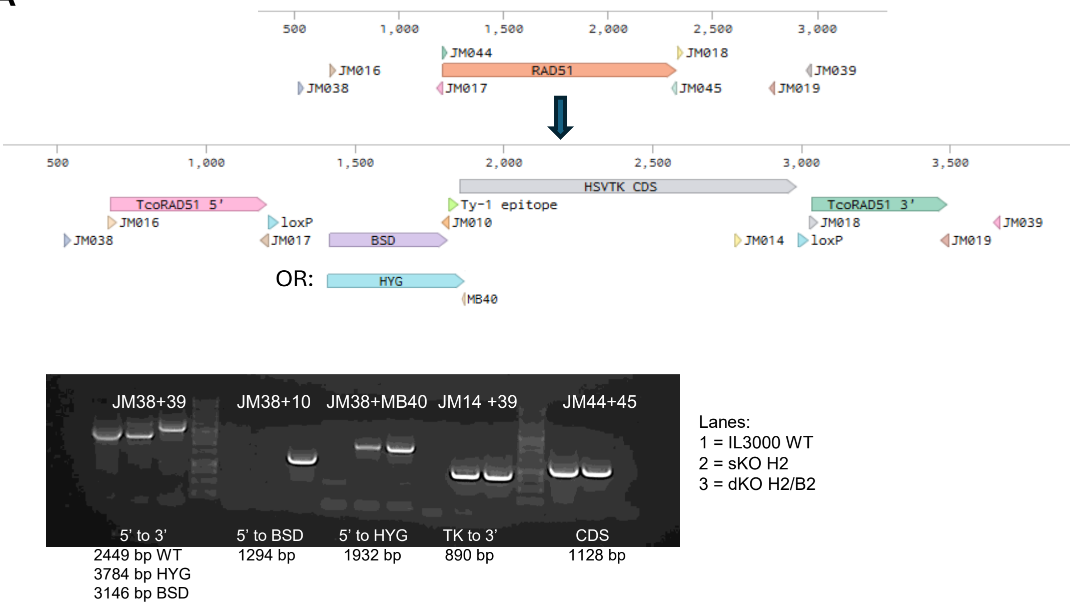


Lanes:
1 = IL3000 WT
2 = sKO H2
3 = dKO H2/B2

**B** TcoBRCA2$^{-/-}$

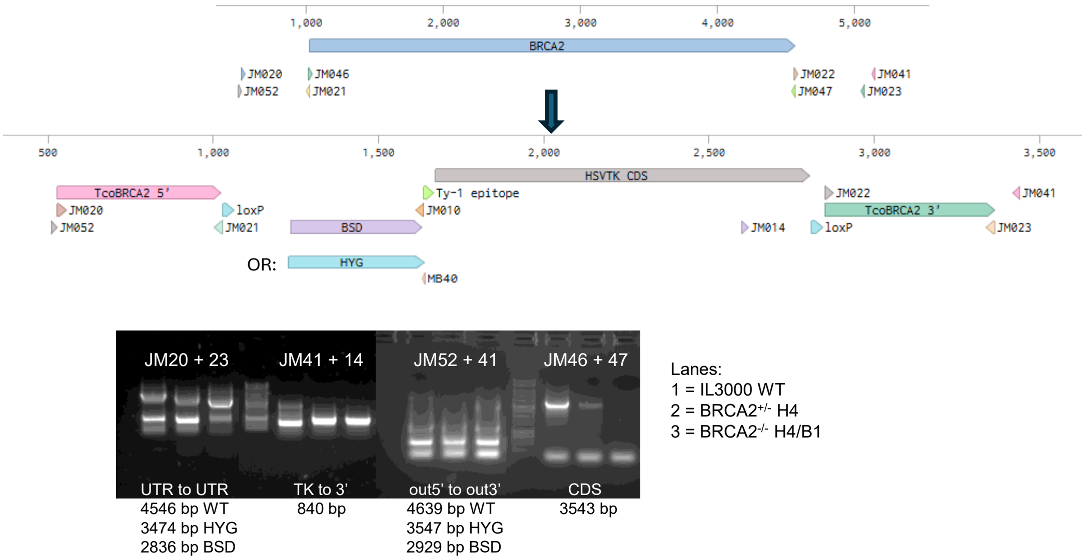


Lanes:
1 = IL3000 WT
2 = BRCA2$^{+/-}$ H4
3 = BRCA2$^{-/-}$ H4/B1

**C**

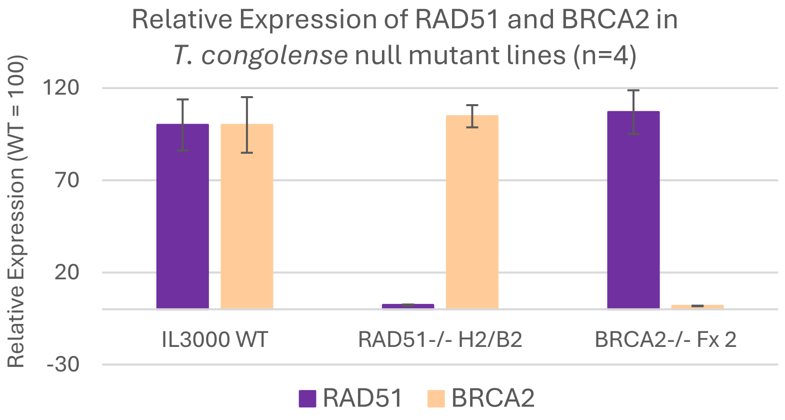

### Figure S2

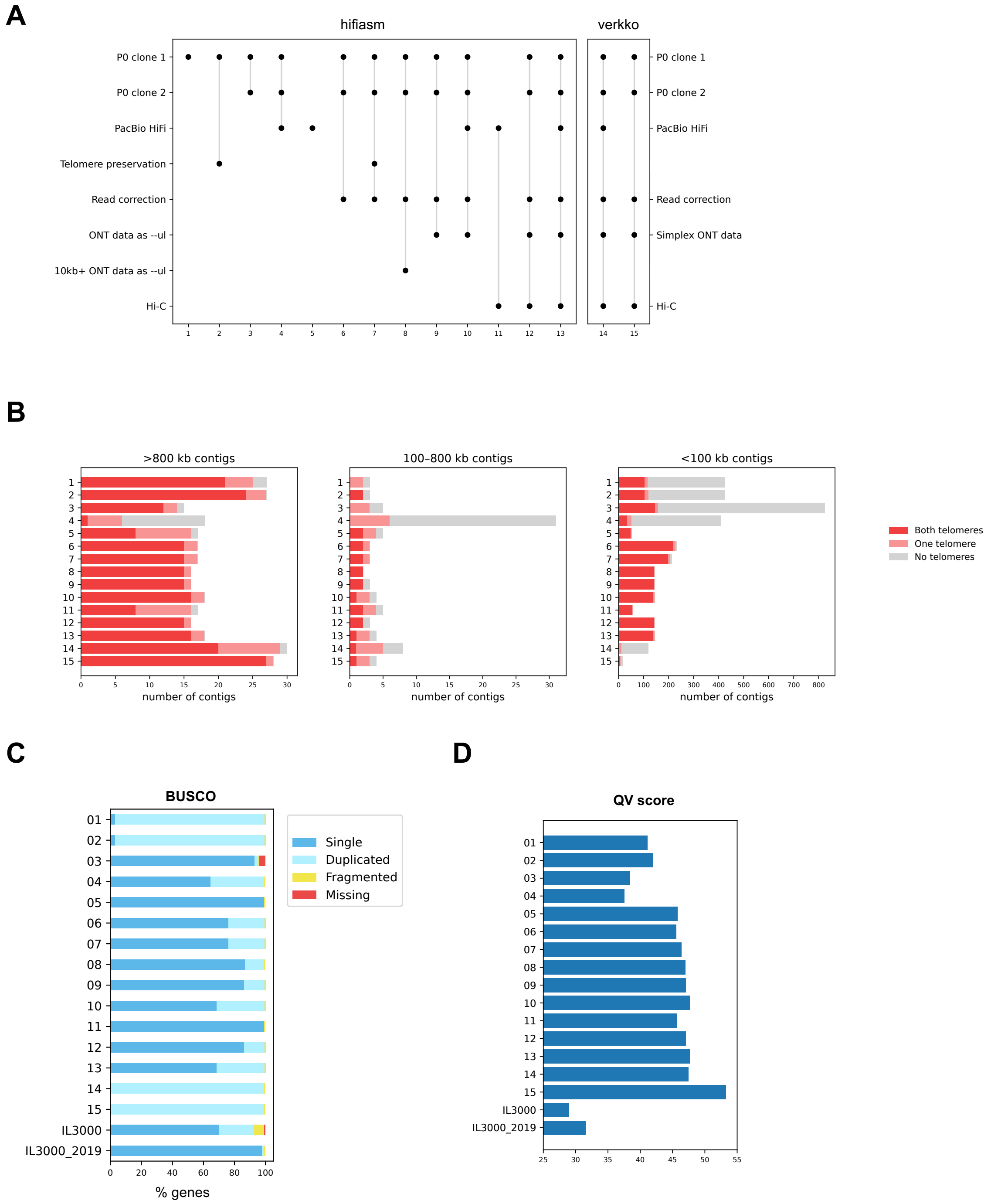
A
hifiasm
verkko
P0 clone 1
P0 clone 2
PacBio HiFi
Telomere preservation
Read correction
ONT data as --ul
10kb+ ONT data as --ul
Hi-C
Simplex ONT data
B
>800 kb contigs
100–800 kb contigs
<100 kb contigs
number of contigs
Both telomeres
One telomere
No telomeres
C
BUSCO
Single
Duplicated
Fragmented
Missing
IL3000
IL3000_2019
% genes
D
QV score
IL3000
IL3000_2019

### Figure S3

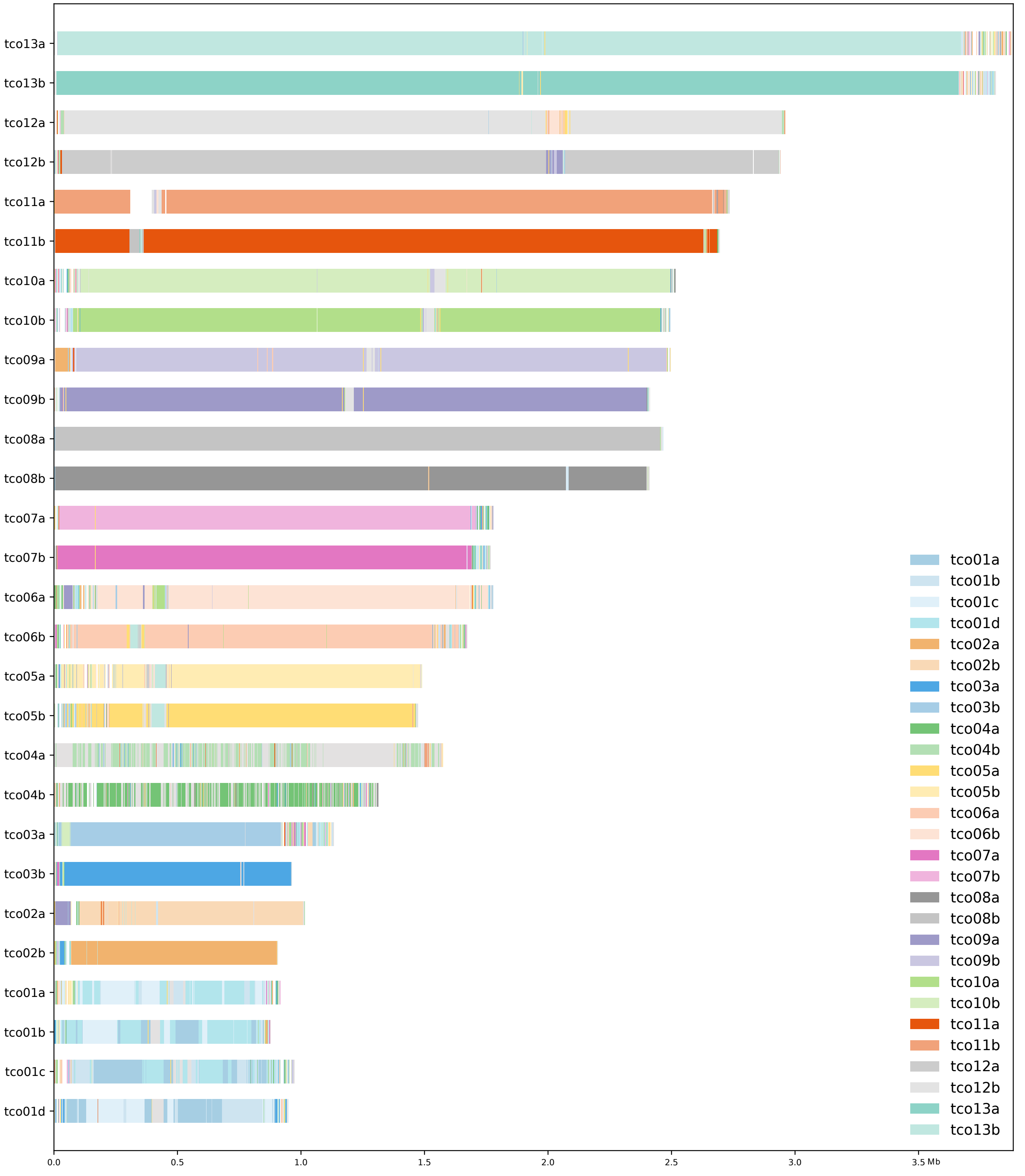

tco13a
tco13b
tco12a
tco12b
tco11a
tco11b
tco10a
tco10b
tco09a
tco09b
tco08a
tco08b
tco07a
tco07b
tco06a
tco06b
tco05a
tco05b
tco04a
tco04b
tco03a
tco03b
tco02a
tco02b
tco01a
tco01b
tco01c
tco01d
0.0
0.5
1.0
1.5
2.0
2.5
3.0
3.5 Mb
tco01a
tco01b
tco01c
tco01d
tco02a
tco02b
tco03a
tco03b
tco04a
tco04b
tco05a
tco05b
tco06a
tco06b
tco07a
tco07b
tco08a
tco08b
tco09a
tco09b
tco10a
tco10b
tco11a
tco11b
tco12a
tco12b
tco13a
tco13b

**Figure S4**

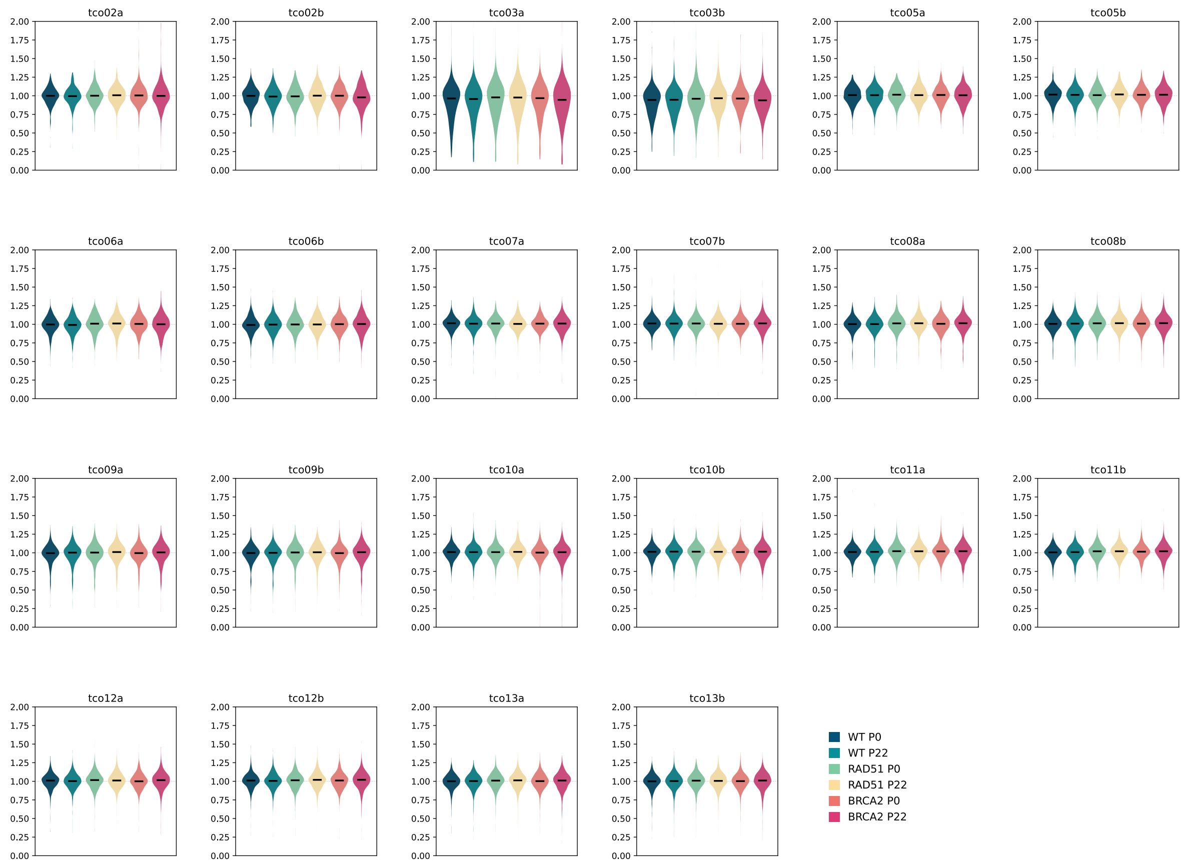
tco02a
tco02b
tco03a
tco03b
tco05a
tco05b
tco06a
tco06b
tco07a
tco07b
tco08a
tco08b
tco09a
tco09b
tco10a
tco10b
tco11a
tco11b
tco12a
tco12b
tco13a
tco13b
0.00
0.25
0.50
0.75
1.00
1.25
1.50
1.75
2.00
WT P0
WT P22
RAD51 P0
RAD51 P22
BRCA2 P0
BRCA2 P22

**Figure S5**

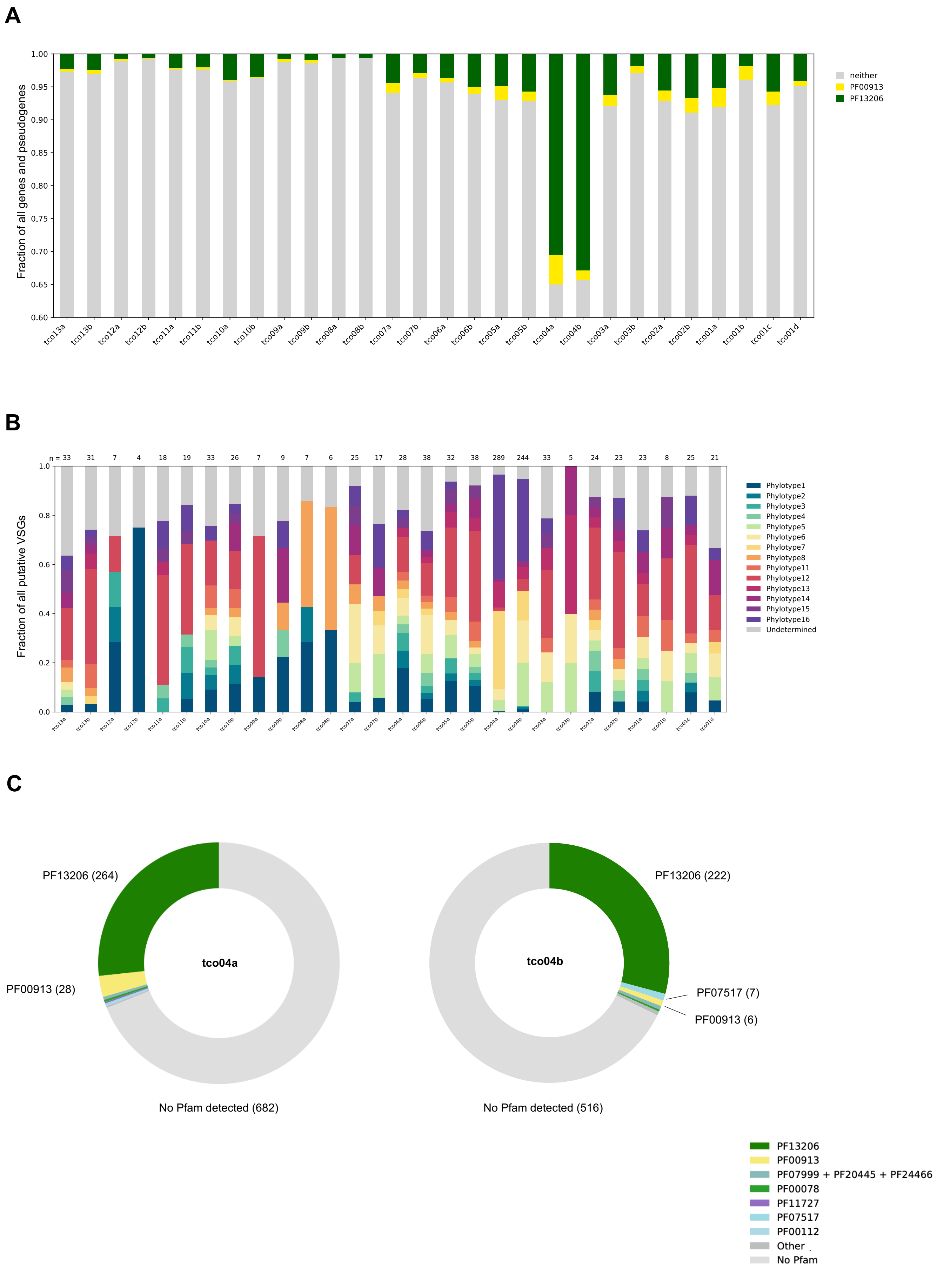
A
Fraction of all genes and pseudogenes
neither
PF00913
PF13206
tco13a
tco13b
tco12a
tco12b
tco11a
tco11b
tco10a
tco10b
tco09a
tco09b
tco08a
tco08b
tco07a
tco07b
tco06a
tco06b
tco05a
tco05b
tco04a
tco04b
tco03a
tco03b
tco02a
tco02b
tco01a
tco01b
tco01c
tco01d
B
n = 33 31 7 4 18 19 33 26 7 9 7 6 25 17 28 38 32 38 289 244 33 5 24 23 23 8 25 21
Fraction of all putative VSGs
Phylotype1
Phylotype2
Phylotype3
Phylotype4
Phylotype5
Phylotype6
Phylotype7
Phylotype8
Phylotype11
Phylotype12
Phylotype13
Phylotype14
Phylotype15
Phylotype16
Undetermined
C
PF13206 (264)
PF00913 (28)
tco04a
No Pfam detected (682)
PF13206 (222)
PF07517 (7)
PF00913 (6)
tco04b
No Pfam detected (516)
PF13206
PF00913
PF07999 + PF20445 + PF24466
PF00078
PF11727
PF07517
PF00112
Other
No Pfam

### Figure S6

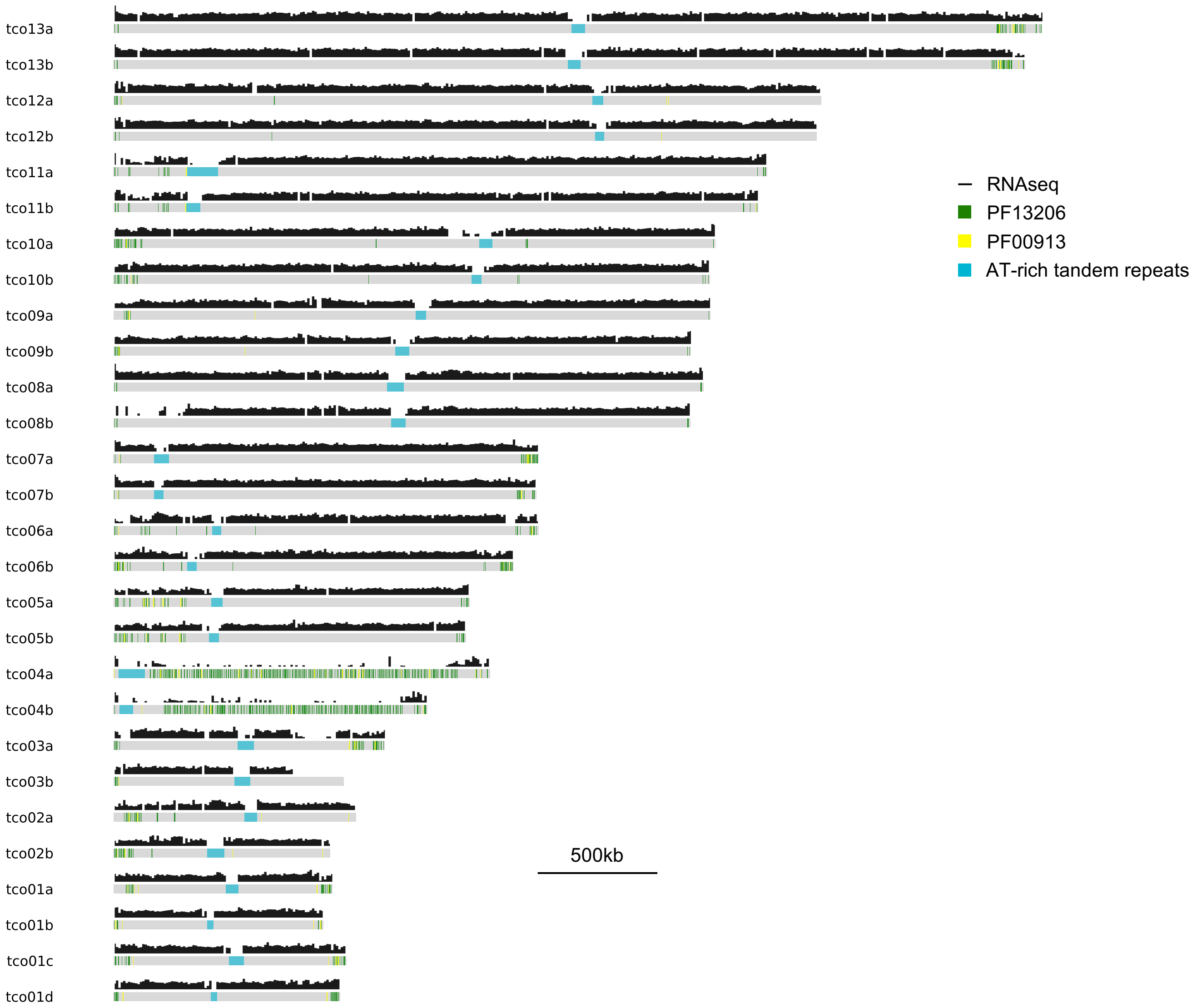
tco13a
tco13b
tco12a
tco12b
tco11a
tco11b
tco10a
tco10b
tco09a
tco09b
tco08a
tco08b
tco07a
tco07b
tco06a
tco06b
tco05a
tco05b
tco04a
tco04b
tco03a
tco03b
tco02a
tco02b
tco01a
tco01b
tco01c
tco01d
RNAseq
PF13206
PF00913
AT-rich tandem repeats
500kb

### Figure S7

**A**

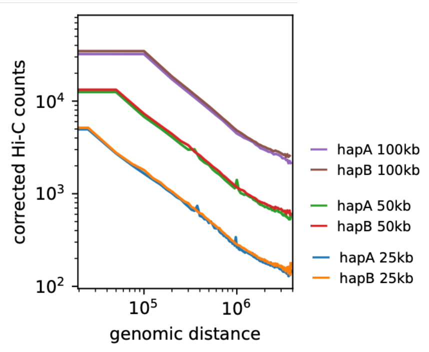


**B**

Pearson correlation matrices
25kb resolution

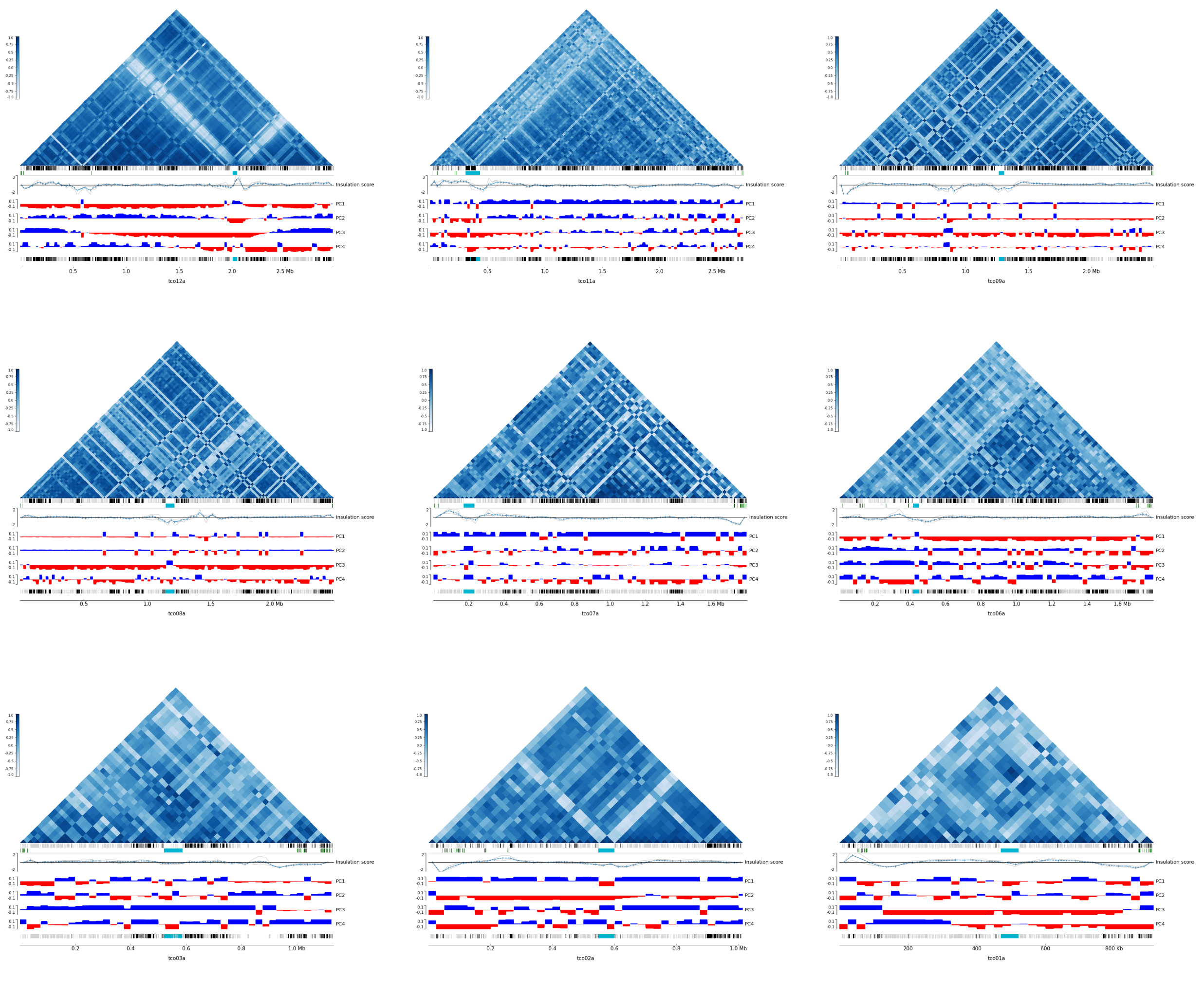


VSGs
AT-rich tandem repeats
genes (+)
genes (-)

#### Figure S8

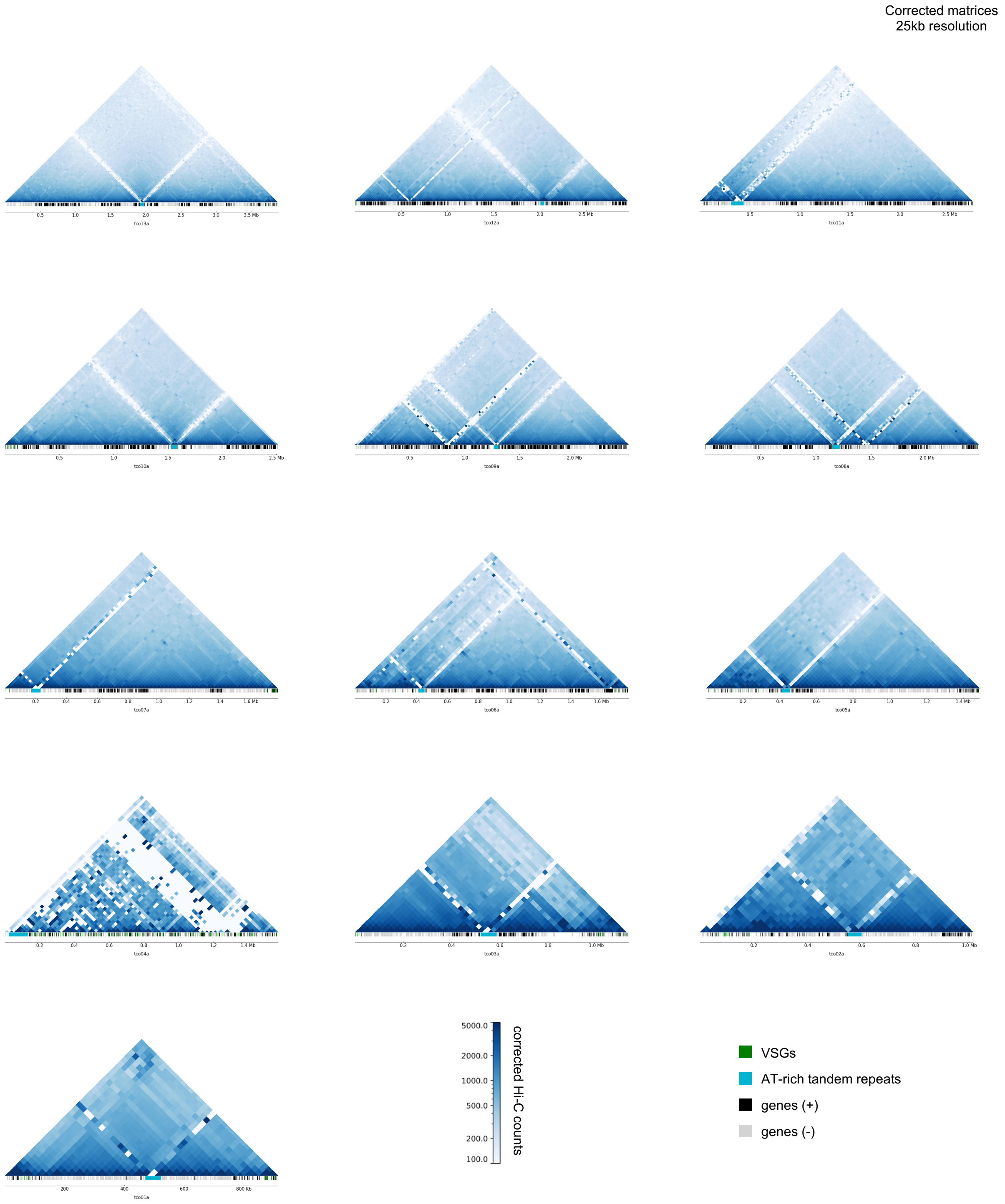
Corrected matrices
25kb resolution
tco13a
tco12a
tco11a
tco10a
tco09a
tco08a
tco07a
tco06a
tco05a
tco04a
tco03a
tco02a
tco01a
corrected Hi-C counts
5000.0
2000.0
1000.0
500.0
200.0
100.0
VSGs
AT-rich tandem repeats
genes (+)
genes (-)

#### Table S1

| | IL3000 | IL3000_2019 | 2026 |
|---|---|---|---|
| # contigs (>= 0 bp) | 2839 | 375 | 189 |
| # contigs (>= 1000 bp) | 2834 | 375 | 189 |
| # contigs (>= 5000 bp) | 1289 | 359 | 189 |
| # contigs (>= 10000 bp) | 503 | 312 | 189 |
| # contigs (>= 25000 bp) | 77 | 185 | 188 |
| # contigs (>= 50000 bp) | 16 | 113 | 44 |
| Total length (>= 0 bp) | 41372041 | 39170526 | 61650343 |
| Total length (>= 1000 bp) | 41368191 | 39170526 | 61650343 |
| Total length (>= 5000 bp) | 36540844 | 39113032 | 61650343 |
| Total length (>= 10000 bp) | 30997589 | 38767969 | 61650343 |
| Total length (>= 25000 bp) | 24679909 | 36629074 | 61636704 |
| Total length (>= 50000 bp) | 22805545 | 34025332 | 56387238 |
| # contigs | 2839 | 375 | 189 |
| Largest contig | 4815855 | 4601516 | 3883400 |
| Total length | 41372041 | 39170526 | 61650343 |
| GC (%) | 48.18 | 45.98 | 47.45 |
| N50 | 1222280 | 1070342 | 2412744 |
| N90 | 4531 | 39286 | 192563 |
| auN | 1512764.1 | 1614293.1 | 2064127.1 |
| L50 | 9 | 9 | 11 |
| L90 | 1435 | 141 | 32 |
| # N's per 100 kbp | 17624.70 | 189.21 | 1.62 |
| # predicted genes (unique) | 15046 | 15879 | 20812 |
| # predicted genes (>= 0 bp) | 15869 + 347 part | 16433 + 16 part | 24000 + 1 part |
| # predicted genes (>= 300 bp) | 13532 + 329 part | 14091 + 15 part | 21299 + 1 part |
| # predicted genes (>= 1500 bp) | 4124 + 117 part | 3946 + 3 part | 7435 + 0 part |
| # predicted genes (>= 3000 bp) | 1060 + 37 part | 999 + 1 part | 2239 + 0 part |
| # predicted rRNA genes | 73 + 9 part | 134 + 22 part | 256 + 32 part |
